# Social deficits induced by pervasive environmental stressors are prevented by microbial or dopaminergic modulation

**DOI:** 10.1101/2022.02.28.482288

**Authors:** Caroline J. Smith, Danielle N. Rendina, Marcy A. Kingsbury, Karen E. Malacon, Dang M. Nguyen, Jessica J. Tran, Benjamin A. Devlin, Madeline J. Clark, Ravikiran M. Raju, Lauren Burgett, Jason H. Zhang, Murat Cetinbas, Ruslan I. Sadreyev, Kevin Chen, Malvika S. Iyer, Staci D. Bilbo

## Abstract

Environmental toxicant exposure, including air pollution, is increasing worldwide. However, toxicant exposures are not equitably distributed. Rather, low-income and minority communities bear the greatest burden, along with higher levels of psychosocial stress. Both air pollution and maternal stress during pregnancy have been linked to neurodevelopmental disorders such as autism, but biological mechanisms and targets for therapeutic intervention remain poorly understood. We demonstrate that combined prenatal exposure to air pollution (diesel exhaust particles, DEP) and maternal stress (MS) in mice induces social behavior deficits only in male offspring, in line with the male bias in autism. These behavioral deficits are accompanied by changes in microglia and dopaminergic circuits in the brain, along with changes in the structure of the gut epithelium and microbiome. Importantly, DEP/MS-induced social deficits in males are prevented by shifting the gut microbiome by cross-fostering at birth and reversed by chemogenetic activation of the dopamine system.

## Introduction

Air pollution represents a global and ever-increasing burden to human health (Cohen et al., 2017; Southerland et al., 2022). Importantly, there are pervasive socioeconomic disparities in the burden of air pollution exposure. This year, Jbaily et al. 2022, showed that low-income and racial minority communities are exposed to higher levels of air pollution than high socioeconomic status, primarily white, communities. These communities are also subjected to fewer resources and higher levels of psychosocial stress (Earnshaw et al., 2013). This convergence of toxicant and psychosocial burdens makes these populations more susceptible to the adverse health consequences of air pollution exposure (McGuinn et al., 2019).

High levels of air pollution, particularly during development, have been linked to several diseases, including autism spectrum disorder (ASD; Rahman et al., 2022; Carter et al., 2022; Volk et al., 2013). Like air pollution, maternal stress during pregnancy has been associated with an increased risk for having a child with ASD (Roberts et al., 2014; Kinney et al., 2008). ASD is a neurodevelopmental disorder primarily characterized by deficits in social behavior and communication, as well as repetitive and restrictive behaviors. ASD is also characterized by a strong male bias in prevalence (4:1) and males tend to be more sensitive than females to early life challenges (Baio et al., 2018; Klein and Flanagan, 2016). Together, these findings suggest that air pollution and maternal stress exposure during pregnancy may synergize to increase ASD risk, particularly in male offspring.

When examining the neurobiological underpinnings of ASD, many brain regions have been implicated as disrupted in function and/or connectivity with other regions. Among these, the mesolimbic reward circuits of the brain have been strongly implicated in both clinical and preclinical models of ASD (Supekar et al., 2018; Walsh et al. 2018; Polk & Ikuta, 2022). Activity in these circuits is critical to reward, motivation, and social interactions. Optogenetic stimulation of nucleus accumbens (NAc)-projecting dopamine neurons in the ventral tegmental area (VTA) increases social behavior in mice (Gunaydin et al., 2014) and dopaminergic signaling in the NAc mediates social play behavior in rodents (Kopec et al., 2018; Manduca et al., 2016). Microglia, the resident immune cells of the brain, respond potently to air pollution (Bolton et al., 2017; Block et al., 2022) and play a critical role in the developmental organization of neural circuits (Dziabis & Bilbo, 2021). Indeed, we recently showed that microglia phagocytose and eliminate dopamine D1 receptors (D1Rs), within the NAc during adolescence in rats, a function that is critical to the development of social play behavior in males but not females (Kopec et al., 2018). However, little is known regarding the impact of prenatal environmental exposures on microglia within the mesolimbic reward system, or on dopaminergic function within this pathway.

ASD is also increasingly recognized to be a whole-body disorder. In particular, the gut-brain axis has emerged as key to the pathophysiology and potential treatment of ASD (Sherwin et al., 2019). Changes in the composition of the gut microbiome and gastrointestinal symptoms have been documented in both children and adults with ASD and clinical trials have found long-lasting improvements in both gastrointestinal and behavioral symptoms following microbiota transfer therapy with gut microbiota from healthy individuals (Kang et al., 2019; 2020). Evidence suggests that changes in gut microbiota in ASD may be driven by the highly restrictive diets of individuals with ASD (Yap et al., 2021), but regardless of the source of these changes, the gut microbiome represents an important site for therapeutic intervention in the amelioration of behavior impairments in this disorder.

Importantly, several studies suggest that microglial function and activity of the dopamine system are impacted by the composition of the gut microbiome. For instance, germ-free mice, which lack microbes completely, have immature, hyper-ramified microglia (Erny et al., 2015; Thion et al., 2018). In multiple mouse models of ASD, bacterial supplementation with the bacterial species *Lactobacillus reuteri* induced neuroplasticity in VTA dopamine neurons and restored sociability (Buffington et al., 2016; Sgritta et al., 2019). While early studies have shown that air pollution exposure shifts the composition of the gut microbiome in adult mice (van den Brule et al., 2021; Liu et al., 2021), nothing is known about how maternal exposure to air pollution during gestation might impact the gut microbiome of offspring, and how these shifts might impact the development of circuits underlying social behavior.

To examine these questions, our lab has established a paradigm in which mouse dams are exposed to a combination of diesel exhaust particles (DEP) and maternal stress (MS) during pregnancy (Bolton et al., 2013; Block et al., 2022). We report that maternal DEP/MS exposure alters microglial morphology and gene expression and decreases dopaminergic tone in the NAc only in male offspring. Moreover, chemogenetic activation of the dopamine system is sufficient to rescue social behavior deficits following DEP/MS in males. Finally, we show for the first time that prenatal DEP/MS alters the composition of the gut microbiome in male offspring only, and that shifting the colonization of the gut microbiome at birth via a cross-fostering procedure prevents sociability deficits in DEP/MS-exposed males.

## Results

### Prenatal DEP/MS exposure induces social behavior deficits in male offspring only

We characterized social behavior in offspring following prenatal exposure to either combined DEP/MS (diesel exhaust particles and maternal stress) or control (CON) conditions. Multiparous *C57Bl/6J* mouse dams were exposed to DEP (50μg/50μl 0.05% PBS-T) via oropharyngeal administration every three days throughout gestation according to Bolton et al., 2013. To induce maternal stress, these dams were placed on a wire mesh grate with no bedding material and only 2/3 of a cotton nestlet from embryonic day (E) 13.5-17.5 (Bolton et al., 2013; Rice et al., 2008). CON dams were given oropharyngeal instillations of the vehicle (50μl 0.05% PBS-T) and control housing (bedding material and a full cotton nestlet). No group differences were observed in maternal weight gain during pregnancy, litter size, sex ratio within a litter, or in offspring body weight (Extended Fig. 1). For complete litter and animal numbers per comparison, as well as complete statistics, throughout the manuscript, see Supplementary Tables 1 & 2, respectively).

The adolescent period – defined as approximately postnatal day (P) 25-40 in the mouse – is one during which social interactions with peers are of heightened importance. Thus, we conducted our behavioral assays during this period (Fig. 1a). We used a three-chambered social preference testing chamber as previously described (Smith et al., 2015; Fig. 1b), in order to assess both sociability (i.e. the preference to investigate a social stimulus over an inanimate object) and social novelty preference (i.e. the preference to investigate a novel social partner as compared to one that is familiar). In the sociability assay, we found that CON males showed a strong preference for a novel sex-, age-, and treatment-matched social stimulus as compared to an object, whereas DEP/MS males did not (Fig. 1c&d). This effect was male-specific as no such deficit was observed in the DEP/MS females. Similarly, in the social novelty preference test, CON males showed a strong preference for a novel social stimulus over a familiar cage mate, whereas DEP/MS males did not (Fig. 1f&g). To ascertain whether these deficits were driven primarily by one prenatal exposure or the other, we also performed the sociability assay following DEP alone or MS alone in male and female offspring. We found that neither treatment on its own induced social deficits, indicating that the synergism between the two is required for social behavior deficits (Extended Fig. 2). Finally, we observed no treatment effects on marble-burying (as a measure of repetitive behavior; Fig. 1h&i) or on anxiety-like behavior (as assessed in the open field test; Fig. 1j&k), suggesting that the effects of DEP/MS may be specific to the social domain, at least during the adolescent period.

**Figure 1.**
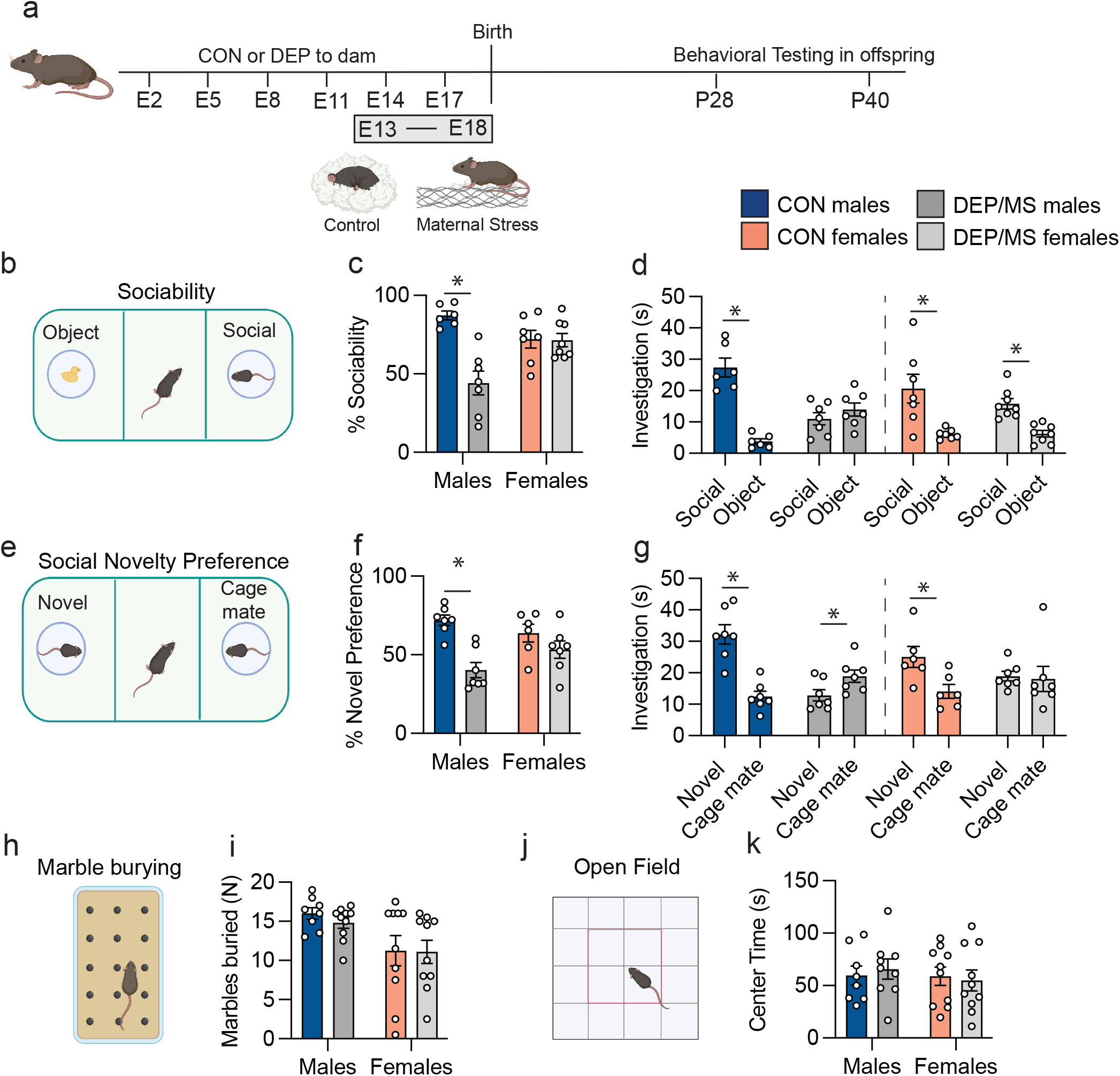
DEP/MS impairs social behavior in male but not female offspring. **a**, Schematic of DEP/MS or CON procedure. **b**, Sociability assay. **c**, Social preference is significantly lower in DEP/MS males as compared to CON males. However, DEP/MS does not alter social preference in females (N=6-8/group; 2-way ANOVA [treatment x sex], treatment: p<0.001, sex: p=0.28, interaction: p<0.001, Bonferonni posthoc: CON male vs. DEP/MS male: p<0.001). **d**, Social investigation time is significantly higher than object investigation time in all groups except DEP/MS males (N=6-8/group, un-paired t-tests (social vs. object): CON M: p<0.001, DEP/MS M: p=0.32, CON F: p<0.01, DEP/MS F: p<0.001). **e**, Social Novelty Preference Test. **f**, Social novelty preference is significantly lower in DEP/MS males as compared to CON males. However, DEP/MS does not alter social novelty preference in females (N=6-7/group; 2-way ANOVA [treatment x sex], treatment: p<0.001, sex: p=0.23, interaction: p=0.04, Bonferonni posthoc: CON male vs. DEP/MS male: p<0.01). **g**, CON males and females significantly prefer a novel social stimuli over a cage mate. DEP/MS males significantly prefer a familiar cage mate, while DEP/MS females show no preference (N=6-8/group, un-paired t-tests (novel vs. cage mate): CON M: p<0.001, DEP/MS M: p=0.04, CON F: p=0.02, DEP/MS F: p=0.84). **h**, Marble-burying. **i**, DEP/MS had no effect on marble burying behavior (N=8-10/group; 2-way ANOVA [treatment x sex], treatment: p=0.63, sex: p=0.004, interaction: p=0.71). **j**, Open Field. **k**, DEP/MS has no effect on center time in the open field (N=8-10/group; 2-way ANOVA [treatment x sex], treatment: p=0.90, sex: p=0.55, interaction: p=0.60). Data represent mean + SEM, *p<0.05. CON: vehicle/control, DEP/MS: diesel exhaust particles/maternal stress. M: males, F: females, E: Embryonic, P: postnatal.

### DEP/MS induces a hyper-ramified, surveillant phenotype in male but not female microglia

Developmental insults, including exposure to environmental chemicals, have been shown to have a particularly potent impact on microglia, the resident immune cells of the brain. During development, microglia play a critical role in the organization of neural circuits via synaptic phagocytosis and elimination, including social circuits (Kopec et al., 2018). Microglia also appear to interact with neurons in other ways, including contacting neurons (Bolton et al., 2017) and ‘trogocytosis’ i.e. synaptic ‘nibbling’ (Weinhard et al., 2018), although these processes remain incompletely understood. Our previous work shows that DEP exposure during gestation increases microglial-neuronal interactions in cortex, and that DEP/MS impairs microglial pruning of thalamocortical synapses in male offspring (Bolton et al., 2017; Block et al., 2022).

Thus, we determined the impact of DEP/MS on microglia in the NAc. We chose to focus on the adolescent/early adult period (P30-60), as this was proximal to our behavioral testing and is a developmental window in which microglia interact with the dopamine system to direct social play behavior (Kopec et al., 2018). Microglial morphology is often taken as an early indicator of alterations in microglial function. Using a MATLAB-based, semi-automatic program for the quantification of microglial ramification (3DMorph; York et al., 2018), we found that NAc-microglia are hyper-ramified following DEP/MS in males but not in females (Fig. 2a). We did not observe any difference in the overall density of microglia between CON and DEP/MS males or females, as assessed using immunohistochemistry for Iba1 quantified by mean pixel intensity (Fig. 2b). To define changes in microglial morphology in males in more detail, we used Imaris 3D image reconstruction software. 3D-reconstruction revealed that the total volume of NAc-microglia was increased following DEP/MS in males (Fig. 2c). Sholl analysis revealed significantly more branch endpoints in DEP/MS male microglia as compared to CON (Fig. 2d), as well as significantly more Sholl intersections (Fig. 2e&f). These findings demonstrate, using multiple approaches, that male microglia are larger and hyper-ramified in the NAc following DEP/MS during the adolescent period.

**Figure 2.**
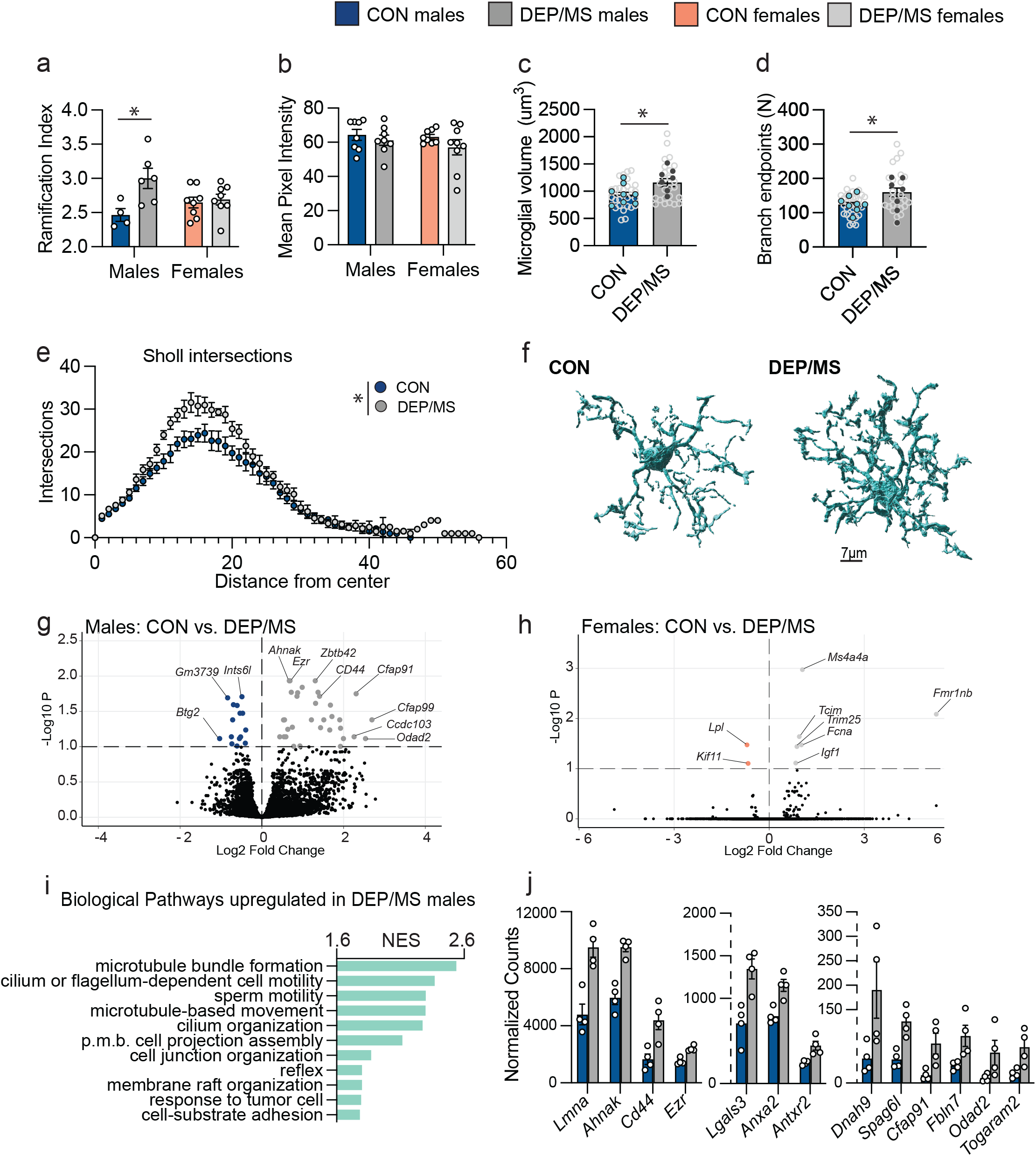
DEP/MS induces a hyper-ramified phenotype in male but not female microglia. **a**, Male, but not female, microglia are hyper-ramified following DEP/MS, as assessed using 3DMorph (N=4-8 animals/group, avg. 11 microglia analyzed/mouse, 2-way ANOVA [treatment x sex], treatment: p=0.012, sex: p=0.55, interaction: p=0.03, Bonferonni posthoc: CON vs DEP/MS males: p=0.02). **b**, No differences were observed between CON and DEP/MS in mean pixel intensity (N=8-9 animals/group, 2-way ANOVA [treatment x sex], treatment: p=0.17, sex: p=0.43, interaction: p=0.65) **c**, Imaris 3D reconstruction revealed larger microglial volume following DEP/MS in males, as well as more branchpoints (**d**) and more sholl intersections (**e**)(N=25-30 microglia from 7-8 animals, nested t-tests [CON vs DEP/MS], volume: p=0.02, branch endpoints: p=0.03, sholl intersections: p=0.02). In **c** and **d**, pale grey dots represent individual microglia while darker dots represent animal averages. **f**, Representative reconstructions of CON and DEP/MS microglia, scale=7μm. **g**, Volcano plots of microglia gene expression reveal numerous genes that are differentially expressed in males following DEP/MS at an adjusted p value cutoff of 0.1 (N=4/group, positive Log2 fold change indicates higher expression following DEP/MS. **h**, In contrast, very few genes are differentially expressed following DEP/MS in females at the same cutoff (N=4/group, positive Log2 fold change indicates higher expression following DEP/MS). **i**, GSEA analysis demonstrates up-regulation in pathways related to cell projection assembly and motility in males following DEP/MS (N=4/group, all pathways reached significance with FDR p<0.05). **j**, Genes related to cell motility and process extension/remodeling were up-regulated in males following DEP/MS. Data represent mean +/- SEM in **a-e** & **j**, *p<0.05. CON:vehicle/control, DEP/MS: diesel exhaust particles/maternal stress, N: number, NES: normalized enrichment score, p.m.b.: plasma membrane bound, GSEA: gene set enrichement analysis.

To gain a deeper understanding of how DEP/MS affects microglial function in sex-specific ways within the NAc, we performed bulk RNA sequencing of microglia isolated from the NAc. In line with our male-biased phenotypes thus far, we found more differentially expressed genes (DEGs) in male microglia following DEP/MS (49; Fig. 2g) than in female microglia (8; Fig. 2h). Notably, more genes (34) were significantly upregulated following DEP/MS in males than were downregulated (15). In keeping with the hyper-ramified morphology of DEP/MS male microglia, gene set enrichment analysis (GSEA) revealed enrichment in male microglia of biological pathways involved in cell motility, extracellular matrix interactions, and cell remodeling/projection assembly (Fig. 2i). Cellular components of similar pathways were also enriched in GSEA analyses (Extended Fig. 3). For example, several of the most highly and differentially transcribed genes encode proteins critical for cell motility/migration (*Ahnak, Dnah9, Odad2, Togaram2, Cfap91, Cfap99*), extracellular matrix interactions and cell adhesion (*CD44, Antxr2, Fbln7, Lgals3*), and intracellular remodeling (*Lmna, Ezr, Syne3, Spag6l*; Fig. 2j). Interestingly, GSEA analysis revealed a distinct set of biological pathways that were enriched in female microglia following DEP/MS (Extended Fig. 3), including response to interferon-β, T cell activation, and regulation of the innate immune response. These patterns were not observed in non-microglial cells in males or females (CD11b-population; Extended Fig. 3).

### DEP/MS alters dopamine circuitry in males but not females

Our findings of a male-specific, hyper-ramified microglial phenotype in the NAc following DEP/MS suggested to us that microglia might be interacting more with neuronal populations in the NAc during development, potentially to alter socially relevant neural circuits, and therefore, social behavior. We recently showed that NAc microglia eliminate dopamine D1 receptors (D1Rs) in this brain region during adolescence (with a specific peak at P30) via phagocytosis, and that this D1R elimination is critical to the developmental trajectory of social play (Kopec et al., 2018). The dopamine system, as well as other neuro-modulatory systems including endogenous opioids and oxytocin, have been shown to mediate social motivation by acting within the NAc (Manduca et al., 2016; Smith et al., 2017, Smith et al., 2018). To determine whether the dopamine system within the NAc was altered following DEP/MS, we first collected tissue punches from this region and conducted qPCR for dopamine receptor mRNA (*Drd1* and *Drd2*), as well as opioid receptors (kappa opioid receptor: *Oprk1*, mu opioid receptor: *Oprm1*) and oxytocin receptor (*Oxtr*). We found that both *Drd1* and *Drd2* mRNA were decreased in the NAc of adolescent male offspring following DEP/MS exposure (Fig. 3a), but not in females (Fig. 3b). mRNA was not decreased for *Oprm1* or *Oxtr* although, interestingly, *Oprk1* mRNA was lower in males and higher in females (Fig. 3a&b). These changes were relatively specific to the NAc, as no changes in these receptors were observed in the amygdala (Extended Fig. 4) – which is also a critical regulator of social behavior via connections with the NAc (Folkes et al., 2020).

**Figure 3.**
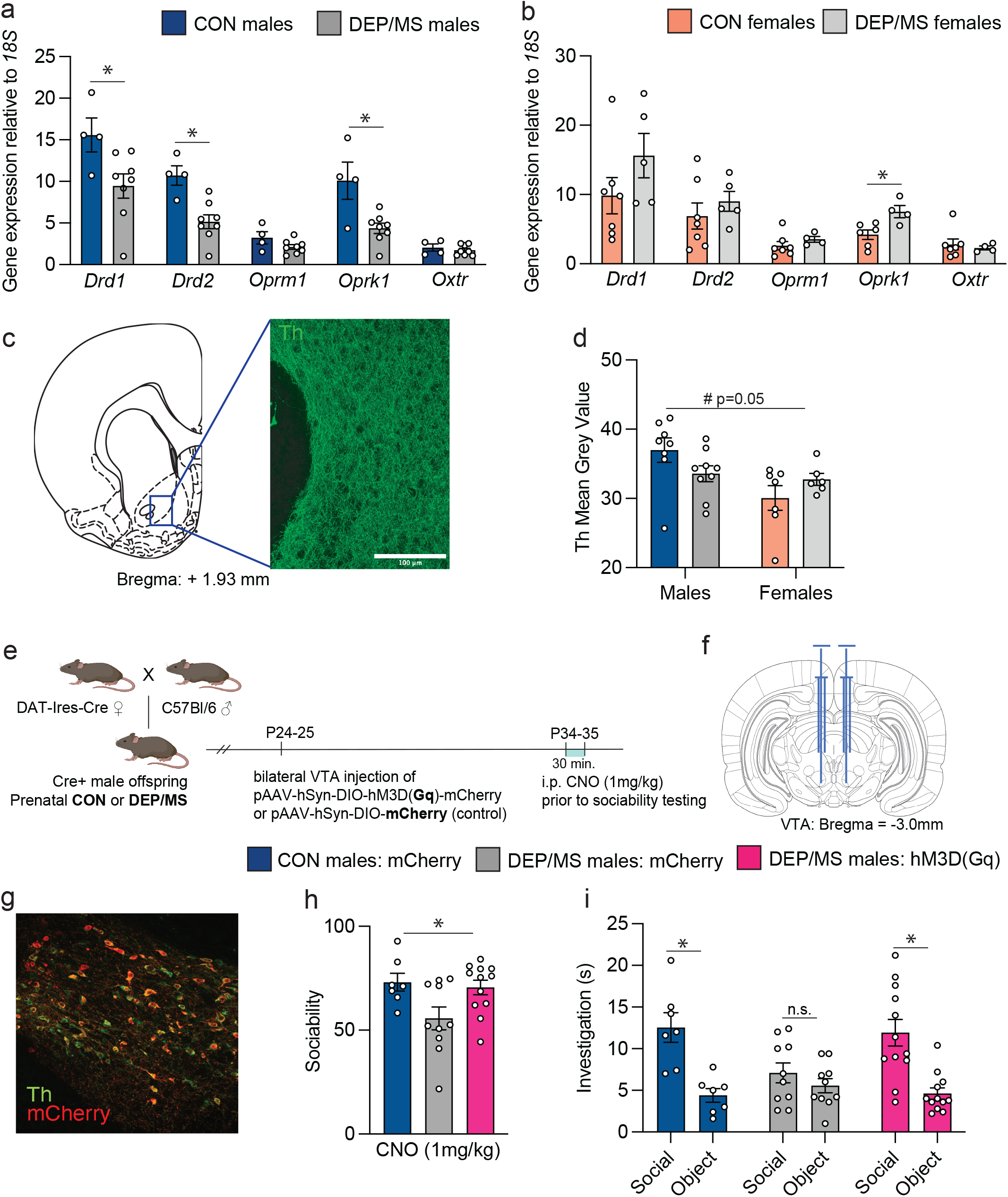
Chemogenetic activation of VTA dopamine neurons rescues sociability in DEP/MS males. **a**, Gene expression in the NAc at P45 for socially relevant receptors in males. mRNA for the dopamine D1 receptor (*Drd1*), D2 receptor (*Drd2*) and the kappa opioid receptor (*Oprk1*) are reduced following DEP/MS in males, while mu opioid receptor (*Oprm1*) and oxytocin receptor (*Oxtr*) mRNA are not (N=4-8/group, unpaired t-tests [CON vs DEP/MS], *Drd1*: p=0.036, *Drd2*: p=0.003, *Oprm1*: p=0.12, *Oprk1*: p=0.008, *Oxtr*: p=0.48). **b**, DEP/MS increases *Oprk1* mRNA in females as compared to CON, but not mRNA for *Drd1, Drd2, Oprm1*, or *Oxtr* (N=4-7/group, unpaired t-tests [CON vs DEP/MS], *Drd1*: p=0.19, *Drd2*: p=0.43, *Oprm1*: p=0.32, *Oprk1*: p=0.017, *Oxtr*. p=0.66). **c**, Representative images of tyrosine hydroxylase (Th) fiber density in the NAc. **d**) Trend towards a significant interaction effect for Th mean grey value in the NAc with lower signal in DEP/MS males as compared to CON, but higher in females at P45 (N=4-9/group, 2-way ANOVA [treatment x sex], treatment: p=0.81, sex: p=0.016, interaction: p=0.053). **e**, Experimental timeline: Following prenatal DEP/MS, Dat-Cre+ male CON and DEP/MS males underwent stereotaxic microinjection of DREADD virus (Gq [excitatory] or mCherry [control] at PND23-24. At P33-35, social behavior was tested 30 min. after i.p. CNO injection. **f**, Mouse brain atlas image of location of microinjection. **g**, Representative 20x image of dopamine fibers (Th) co-labeled with mCherry in the VTA. **h**) Social preference is significantly increased in DEP/MS males following chemogenetic activation of dopaminergic cells in the VTA (N=7-12/group, one-way ANOVA [treatment], p=0.025). **i**, Chemogenetic activation of dopaminergic cells in the VTA restores social preference (N=7-12/group, unpaired t-test [social vs. object], p<0.001. Data are represented as mean + SEM, *p<0.05, #p=0.05. VTA: ventral tegmental area, NAc: nucleus accumbens, Th: tyrosine hydroxylase, P: postnatal, CON: vehicle/control, DEP/MS: diesel exhaust particles/maternal stress, i.p.: intraperitoneal, CNO: Clozapine-N-oxide, DREADD: Designer Receptors Exclusively Activated by Designer Drugs.

Given the decrease in dopamine receptor expression in males, we next asked whether this might be due to increased microglial phagocytosis of D1R at P30. However, we observed no differences in microglial engulfment of D1R using IMARIS 3D volumetric reconstructions of D1R and microglia (Iba1; Extended Fig. 5). An alternative hypothesis is that reduced dopaminergic input from the VTA to the NAc might account for the lower expression of NAc-dopamine receptors following DEP/MS in males. To assess whether these changes in receptor expression were matched by changes in such dopaminergic input to the NAc, we used immunohistochemistry to quantify tyrosine hydroxylase (Th) fiber density within the NAc. We observed a significant interaction effect whereby Th mean grey value (a measure of density) was decreased in males but increased in females following DEP/MS (Fig. 3c&d). These data suggest that the decrease in dopamine receptor expression in DEP/MS males is driven by decreased dopamine input from the VTA into the NAc.

### Chemogenetic activation of the dopamine system rescues male social deficits following DEP/MS

Based on the reduction in Th-fiber density in the NAc in male offspring following DEP/MS, we asked whether chemogenetic activation of the dopamine system would rescue social deficits in male adolescent offspring following DEP/MS. A Dat-Ires-Cre mouse line was used to generate offspring expressing Cre under the control of the dopamine transporter 1 (DAT) promotor (Fig. 3e). Cre+ offspring were exposed to either CON or DEP/MS prenatally. On P24-25, male offspring from both treatment conditions underwent stereotaxic surgery to transfect VTA neurons with either a control Cre-dependent virus expressing an mCherry fluorescent reporter (pAAV-hSyn-DIO-mCherry) or a virus encoding the excitatory DREADD receptor hM3D(Gq) (pAAV-hSyn-DIO-hM3D(Gq)-mCherry; Fig. 3f). Following 10 days for recovery from surgery and to allow for complete transfection of the VTA (Fig. 3g), male offspring underwent sociability testing. Subjects were injected intraperitoneally (I.P.) with clozapine-N-oxide (CNO; 1mg/kg) to activate the excitatory DREADD receptors 30 min prior to testing on the sociability assay. We found that CON males transfected with the control virus showed a significant preference for the social stimulus, as we have previously observed (Fig. 3h). As predicted, DEP/MS males transfected with the control virus displayed no social preference (Fig. 3h). In contrast, DEP/MS males transfected with the excitatory DREADD receptor showed a reinstatement of their preference for a social stimulus following CNO administration (Fig. 3h). Both CON males transfected with a control virus and DEP/MS males transfected with the excitatory DREADD virus spent significantly more time investigating the social stimulus as compared to an object, while DEP/MS males transfected with the control virus spent equal amounts of time investigating the animal and the object (Fig. 3i). Together, these findings suggest that activating VTA-dopamine neurons is sufficient to restore sociability following prenatal DEP/MS in males.

### DEP/MS shifts the composition of the gut microbiome and epithelium

Our results so far demonstrate that prenatal exposure to DEP/MS leads to male-specific changes in social behavior that are accompanied by microglial hyper-ramification and decreased dopaminergic tone within the NAc. Importantly, changes in all three of these endpoints have previously been observed following modulation of the gut microbiome. For instance, changes in the composition of the gut microbiome are common in patients with ASD and microbiota transfer therapy with gut microbiota from healthy controls has been efficacious for the amelioration of social behavior impairments in ASD (Kang et al., 2019; 2020). Similarly, in animal models of ASD such as maternal high-fat diet and Shank 3b knockout (KO) mice, supplementation of the gut microbiome with *L. reuteri* rescues social behavior deficits by modulating activity of the dopamine system (Buffington et al., 2016; Sgritta et al., 2019). Microglia are also exquisitely sensitive to the composition of the gut microbiome (Erny et al., 2015; Thion et al., 2018). Indeed, microglial hyper-ramification, very similar to what we observe in DEP/MS males, has been reported in germ-free mice (Erny et al., 2015).

Based on these findings, we next asked whether DEP/MS exposure impacted the composition of the gut microbiome in offspring. We used bacterial 16S sequencing from cecal contents in male and female offspring during early adulthood (P45: following the end of behavioral testing) to assess the community structure of the gut microbiome in male and female offspring. Measures of alpha diversity quantify overall community richness (how many bacterial taxa are present) or evenness (how evenly abundant the taxa are that form the community) within the gut microbiome of an individual animal. We found that community evenness was significantly increased in DEP/MS males as compared to control males (Pielou’s evenness; Fig. 4a). Principal Coordinate Analysis (PCoA) of quantitative beta diversity indices, which evaluate microbial community structure between individuals, revealed distinct clustering of microbiome profiles in CON and DEP/MS males (Fig. 4b). Permutational multivariate analysis of variance (PERMANOVA) analysis revealed significant differences in the phylogenetic relatedness and abundance of microbial communities between CON and DEP/MS males (Bray-Curtis dissimilarity and weighted UniFrac). Differences were not evident at the level of individual taxa. Notably, no changes in either alpha or beta diversity were found in female offspring (Fig. 4c&d).

**Figure 4.**
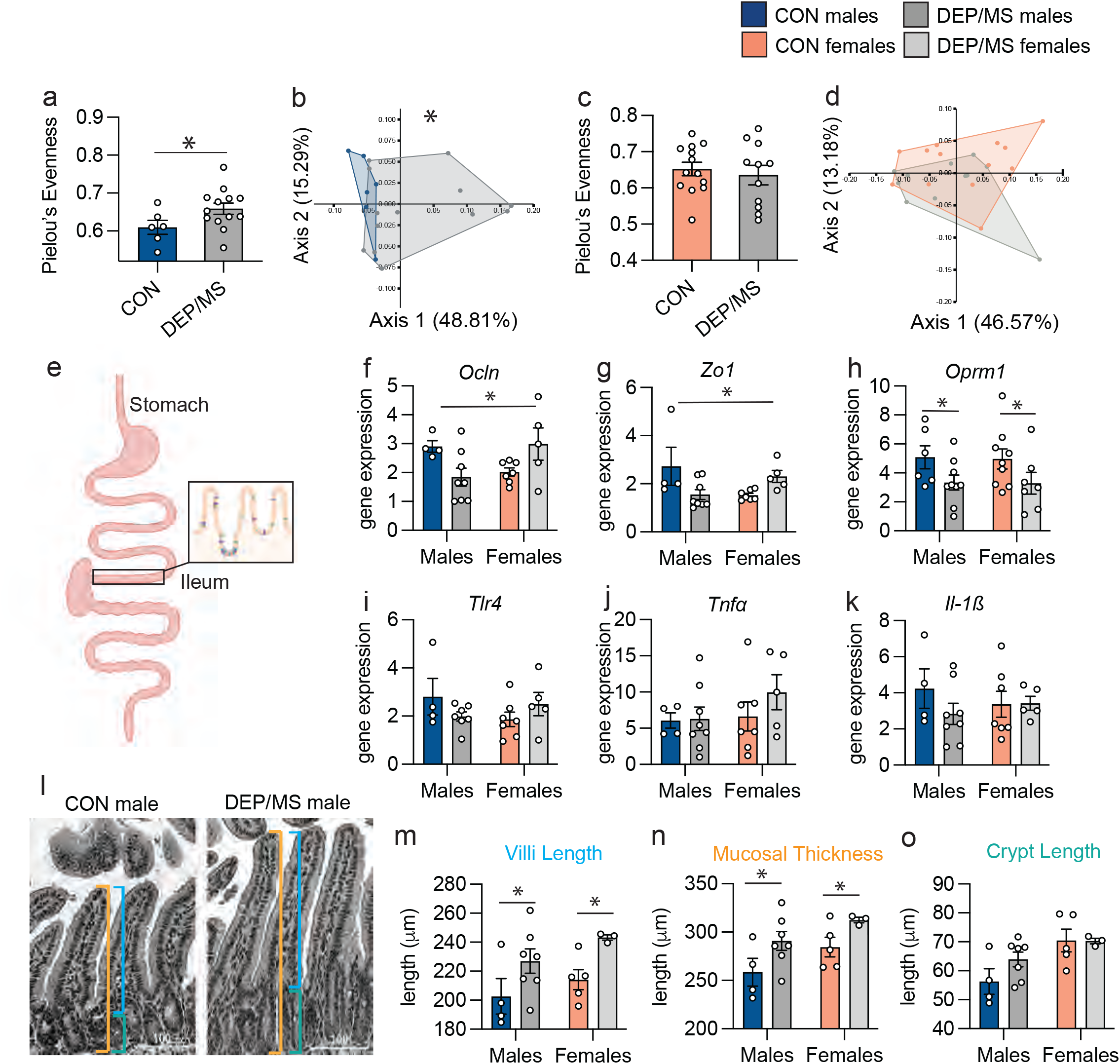
DEP/MS shifts the composition of the gut microbiome and epithelium. **a**, Pielou’s evenness was significantly higher in DEP/MS males as compared to CON (N=6-13/group, p=0.02). **b**, CON and DEP/MS males clustered distinctly in PCoA analyses (N=6-13/group, weighted UniFrac dissimilarity, p=0.03). **c-d**, No changes in alpha or beta diversity were observed in females following DEP/MS exposure (N=11-13/group, Pielou’s evenness: p=0.66, weighted UniFrac dissimilarity: p=0.29). **e**, Representative image of section of the ileum assessed. **f-g**, DEP/MS exposure decreases tight-junction mRNA (*Ocln*, **f**; *Zo1*, **g**) in males, but increases expression in females (N=4-8/group, 2-way ANOVA [treatment x sex], *Ocln*: interaction: p<0.01, *Zo1*: interaction: p<0.01). **h**, DEP/MS decreases *Oprm1* expression in both males and females as compared to CON (N=6-9/group, 2-way ANOVA [treatment x sex], treatment: p=0.02). **i-k**, DEP/MS has no effect on proinflammatory gene expression in the ileum (*Tlr4*, i; *TNFα*, j; *Il-1β*, N=4-8/group, 2-way ANOVA [treatment x sex], *Tlr4*: treatment: p=0.82, *TNFα*: treatment: p=0.38, *Il-1β*: treatment: p=0.36). **l**, Representative H&E staining of epithelium within the ileum. lines: orange=mucosal thickness, blue= villus length, grey=crypt length. Scale=100μm. **m**, DEP/MS increases villi length and mucosal thickness (**n**) in both males and females as compared to CON but has no effect on crypt length (**o**; N=4-7 animals/group, 2-way ANOVA [treatment x sex], villi length: treatment: p=0.01, mucosal thickness: treatment: p=0.01, crypt length: treatment: p=0.24). Data represent mean + SEM, *p<0.05. CON: control, DEP/MS: diesel exhaust particles/maternal stress.

Microbes within the gut interface directly with the intestinal epithelium and are important determinants of epithelial structure and immunity. The tight-junction proteins Occludin (*Ocln*) and Zonula occludins-1 (*Zo1*) stabilize the gap junctions between intestinal epithelial cells, preventing the translocation of bacteria and/or their metabolites from the intestinal lumen into the body. Individuals with ASD often report comorbid gastrointestinal dysfunction and evidence for disruption of the gut epithelial barrier – including changes in *Ocln* and *Zo1* expression – has been found in patients with ASD (Lefter et al., 2019, Nalbant et al., 2021). Therefore, we also assessed gene expression for these tight junction proteins within the ileum, the segment of the small intestine proximal to the cecum (Fig. 4e). We observed a sex-specific effect of DEP/MS (decreased in males but increased in females) on *Ocln* (Fig. 4f) and *Zo1* (Fig. 4g) mRNA. A similar pattern of *Ocln* and *Zo1* expression was also observed in the duodenum in males, but not in the colon, suggesting that changes in tight-junction proteins are specific to the small intestine (Extended Fig. 6). Constipation and diarrhea are the predominant components of GI dysfunction in ASD (Lefter et al., 2019). We found that mRNA transcript for the mu (μ) opioid receptor (*Oprm1*) – which is a critical regulator of gut motility - was decreased in both males and females following DEP/MS (Fig. 4h). Interestingly, we observed no changes in the proinflammatory genes *Tlr4, Tnfα*, or *Il-1β* in either the small (Fig. 4i-k) or large (Extended Fig. 6) intestine, suggesting that these effects are not due to current inflammation, per se.

The entire structure of the intestinal epithelium itself is also sensitive to gut microbial composition. For example, germ-free mice exhibit poorly developed villi and shallow crypts within the lamina propria (which sits directly below the epithelial cell layer; Abrams et al., 1963) as well as slowed turn-over and renewal of epithelial cells and mucus as compared to conventionally housed mice (Arike et al., 2020). Similarly, antibiotic treated mice display reduced proliferative activity of intestinal epithelial cells (Park et al., 2016). Thus, we used hematoxylin/eosin (H&E) staining of the intestinal epithelium in the ileum to assess the impact of DEP/MS on the overall structure of the gut (Fig. 4l). Both villi length (Fig. 4m) and mucosal thickness (Fig. 4n), but not crypt length (Fig. 4o), were increased in both male and female offspring following DEP/MS exposure. Together, these findings demonstrate that the gut microbiome is significantly altered in male offspring only following DEP/MS, and that these changes are accompanied by pervasive changes in the intestinal epithelium – suggestive of decreased gut barrier function specifically in male offspring.

### DEP/MS shifts microglial gene expression towards a germ-free phenotype

The observed changes in the composition of the gut microbiome of male offspring following DEP/MS, as well as microglial hyper-ramification, led us to ask whether there are similarities in microglial gene expression following DEP/MS and other mouse models of microbial disruption. Indeed, the hyper-ramification phenotype we observe is strikingly similar to microglial hyper-ramification that has been observed in germ-free male mice as compared to conventionally housed mice (Erny et al., 2015). To answer this question, we compared gene differentials in male microglia following DEP/MS from our RNA sequencing analysis (see Fig. 2) to microglial genes that are differentially expressed in germ-free adult male mice in a published dataset (Thion et al., 2018). We used stratified Rank-Rank Hypergeometric Overlap (RHHO) analysis (Cahill et al., 2018) to compare concordant and discordant patterns of gene expression between these two datasets. We found a significant concordance between genes altered in DEP/MS and germ-free male microglia, particularly in up-regulated genes (Fig. 5a). To determine whether this concordance was specific, we also assessed the relationship between DEP/MS microglial gene differentials and gene differentials following acute immune activation (2h after lipopolysaccharide: LPS at P60, Hanamsagar et al., 2017; Fig. 5b) and across typical development (Hanamsagar et al., 2017; Fig. 5c). Interestingly, we found significant discordance between DEP/MS microglial transcription and both LPS-induced and developmentally regulated transcription. These findings suggest that genes that are up-regulated following DEP/MS or germ-free conditions in male microglia are down-regulated following immune activation or into adulthood (P4-P60). This is in line with the idea that germ-free microglia are perhaps immune-incompetent and immature, and suggests that a similar phenotype might be induced by DEP/MS exposure in male microglia.

**Figure 5.**
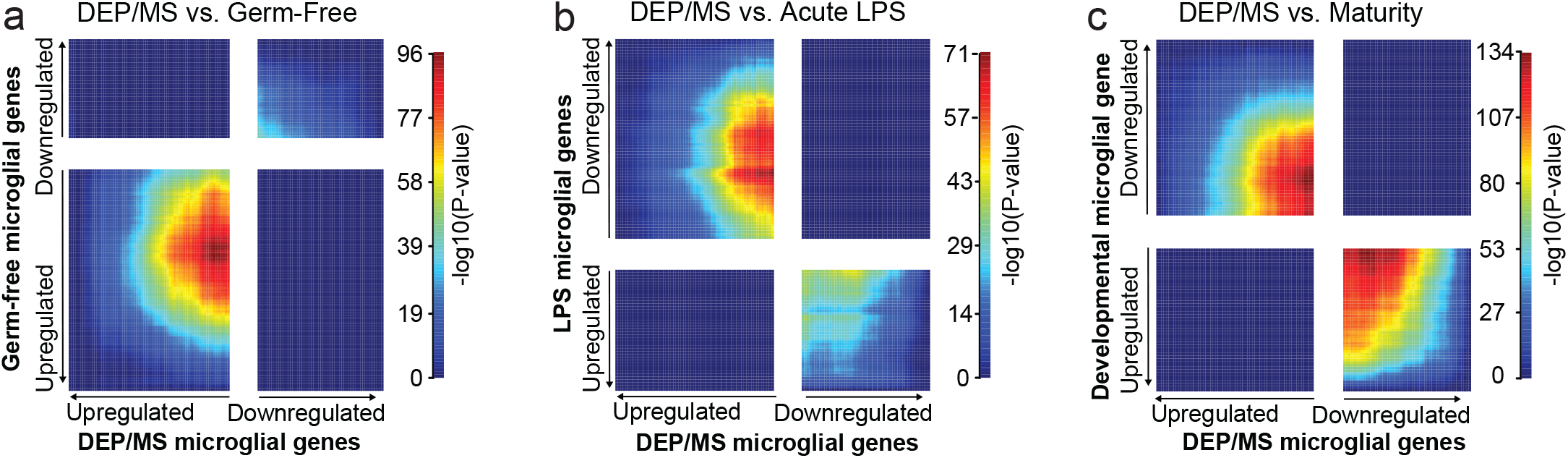
DEP/MS shifts microglial gene expression towards a germ-free phenotype. **a-c**, Stratified Rank-Rank Hypergeometric Overlap (RHHO) analysis on RNA sequencing of isolated microglia from the NAc of male offspring following CON or DEP/MS (N=4/group). Gene rank values were calculated by performing un-paired t-tests between CON and DEP/MS. Average values for each group were then used to calculate the fold-change in mean expression and the sign of this fold-change was then multiplied by the Log10(p value) **a**, DEP/MS-regulated microglia genes were compared to gene differentials in germ-free adult male mice as compared to conventionally housed (Thion et al., 2018). RHHO analysis revealed significant concordance between genes up-regulated in the two datasets. **b**, DEP/MS-regulated microglia genes were compared to gene differentials in lipopolysaccharide (LPS) treated mice as compared to saline control (Hanamsagar et al., 2017). RHHO analysis revealed significant discordance between genes in the two datasets, whereby genes that were up-regulated following DEP/MS were down-regulated following LPS treatment. **c,** DEP/MS-regulated microglia genes were compared to gene differentials across development (P4-P60; Hanamsagar et al., 2017). RHHO analysis revealed significant discordance between expression changes in the two datasets, whereby genes that were up-regulated following DEP/MS were down-regulated with advancing age. DEP/MS: diesel exhaust particles/maternal stress.

### Cross-fostering at birth prevents DEP/MS-induced social deficits in male offspring

Given the well-established link between gut microbiota and social behavior (for review see Sherwin et al., 2019), along with the fact that we observed corresponding male-specific changes in the gut microbiome and social behavior, we next asked whether shifting the gut microbiome towards a CON-typical composition would prevent social behavior deficits in DEP/MS male offspring. Cross-fostering on the day of birth has been shown to reliably shift the composition of the offspring gut microbiome to match that of the foster mother over the birth mother (Daft, et al. 2015; Treichel et al., 2019; Daoust et al., 2021). DEP/MS exposed pups were fostered to either a different DEP/MS dam on the day of birth (D → fD) or to a CON dam (D→ fC). Similarly, CON exposed pups were fostered to either a different CON dam on the day of birth (C → fC) or to a DEP/MS dam (C→ fD). They were then tested on social behavior assays during adolescence prior to sacrifice and sample collection for microbiome and gut analyses (Fig. 6a). Since our fostering manipulation required exposure to a new mother, we characterized maternal behavior to rule out that any changes we observed were simply due to changes in maternal care. Maternal care was assessed three times daily (morning, afternoon, and evening [dark phase]) for the first three days of life. We found no differences between DEP/MS and CON dams in time spent on the nest nursing (Fig. 6b) or in licking and grooming behaviors (Fig. 6c; for further analysis see Extended Fig. 7).

**Figure 6.**
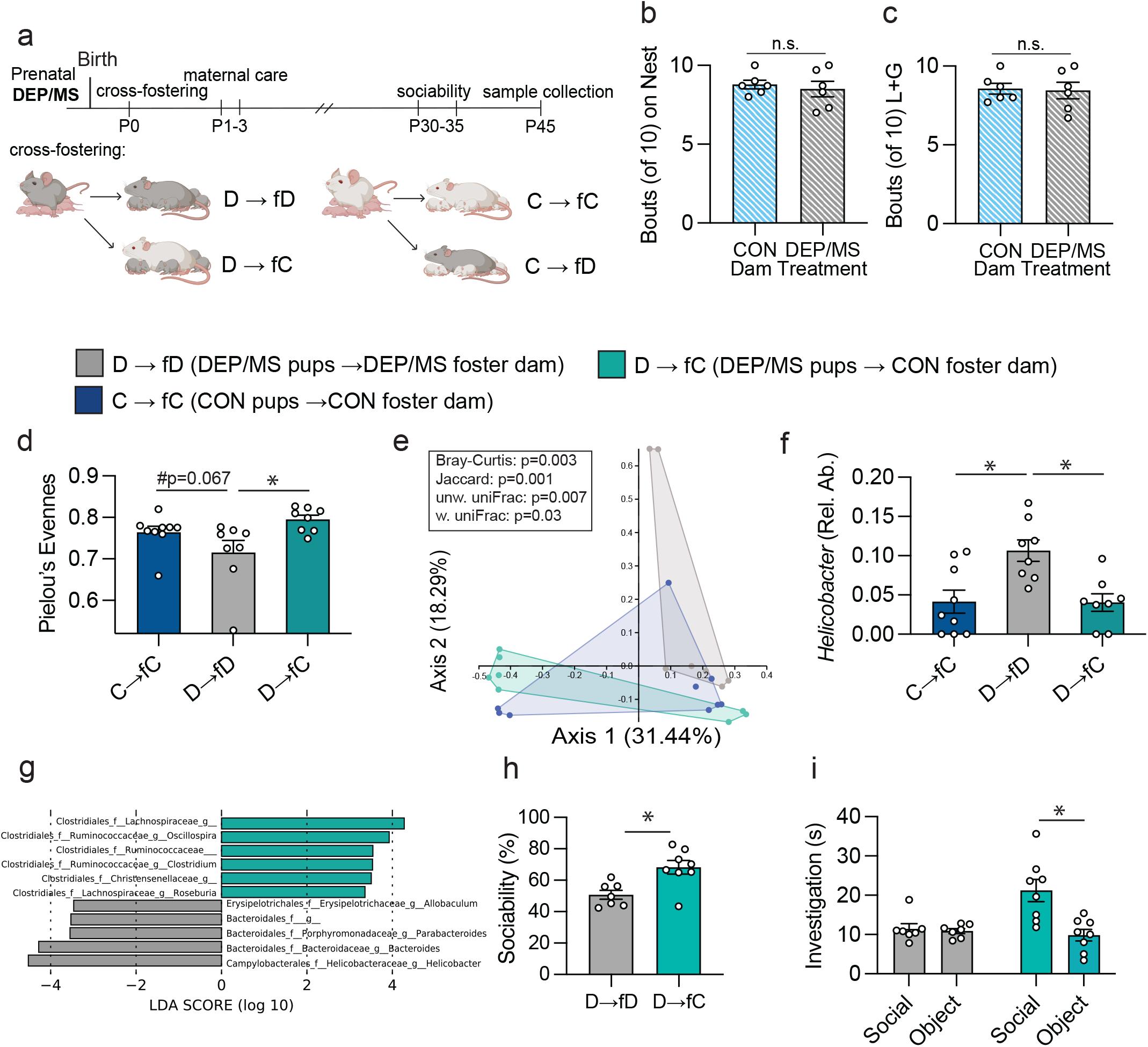
Cross-fostering at birth prevents DEP/MS-induced social deficits in male offspring. **a**, Schematic of cross-fostering procedure and subsequent behavioral testing and sample collection. **b**, Neither time spent on the nest, nor licking and grooming (**c**, L+G= licking and grooming) differed between CON and DEP/MS foster dams (N=6/group, un-paired t-tests [CON vs. DEP/MS]: time on nest: p=0.63, L+G: p=0.86). **d**, Pielou’s evenness differed significantly according to condition (N=7-9/group, p=0.01). D → fC males had significantly higher evenness within the cecal microbiome as compared to D → fD males (p=0.009) while evenness trended towards a difference between C → fC and D → fD males (p=0.067) **e**, The composition of the gut microbiome differed significantly according to condition on all four metrics of beta diversity (N=7-9/group, Bray-Curtis: p=0.003, Jaccard Index: p=0.001, unweighted UniFrac: p=0.007, weighted UniFrac: p=0.03, Bray-Curtis shown in PCoA). **f-g**, LEfSe revealed several genera of bacteria that were differentially abundant in DEP/MS-DEP/MS and DEP/MS-CON males including Helicobacter, Parabacteroides, and Bacteroides (N=7-9/group, 1-way ANOVA [treatment], p=0.002, posthoc: D→fD vs D→fC: p=0.007, D→fD vs C→fC: p=0.006, C→fC vs D→fC: p=0.99). **h**, Fostering to a CON dam on the day of birth significantly increased sociability in male DEP/MS offspring (N=7-8/group, unpaired t-test [D → fD vs. D → fC], p=0.005) and restored a preference for the social stimulus (**i**, (N=7-8/group, unpaired t-tests [Social vs. Object], D → fD: p=0.70, D → fC p=0.003). Data represent Mean +/- SEM, *p<0.05. CON: vehicle and control, DEP/MS: diesel exhaust particles/maternal stress, p: postnatal, w.: weighted, unw.: unweighted, Rel. Ab.: relative abundance.

To verify that cross-fostering of DEP/MS pups to a CON dam would shift the gut microbiome towards a CON-typical phenotype, we utilized 16S sequencing of cecal contents at P45. Indeed, alpha diversity was significantly different overall between the treatment groups (Pielou’s Evenness, Fig. 6d). Pairwise comparisons showed a trend towards lower evenness in D → fD male offspring as compared to C→ fC (p=0.067). D→ fC males had significantly higher evenness than D → fD males (Fig. 6e) that did not differ from C→ fC males. Furthermore, PERMANOVA analysis revealed significantly divergent microbial community structure between D → fD, D→ fC, and C→ fC males in all four beta diversity indices (Fig. 6e, Bray-Curtis, Jaccard dissimilarity, unweighted and weighted UniFrac). In pairwise comparisons, the microbiome of D → fD males differed significantly from that of C→ fC, and D→ fC males. Linear discrimination analysis effect size (LEfSe) identified several genera of bacteria that distinguished the shift within the gut microbiome between D → fD and D→ fC male offspring (Fig. 6f-g). Among these, *Helicobacter, Bacteroides* and *Parabacteroides* were more abundant in D → fD male offspring, while *Lachnospiraceae*, and *Oscillospira* were more abundant in D→ fC male offspring. *Helicobacter pylori (H. pylori*) has been implicated in gut inflammation and can lead to conditions such as acute gastritis (Boukthir et al., 2007; Yu et al., 2014). Both *Parabacteroides* and *Bacteroides* have been shown to be differentially abundant in the gut microbiome in human patients with ASD (Strati et al., 2017; Sharon et al., 2019).

We also asked whether cross-fostering to CON dams would rescue any of the changes we observed in the intestinal epithelium. Interestingly, neither *Ocln* nor *Zo1* mRNA, which were decreased in male offspring following DEP/MS, differed between D → fD and D→ fC males in the small intestine at P45 (Extended Fig. 8). Similarly, villi length and mucosal thickness were not impacted by fostering to a CON dam on the day of birth. However, *Oprm1* mRNA (which was reduced following DEP/MS) tended to be increased in D→ fC males as compared to D → fD males (p=0.08), and crypt length and *Il-1ß* mRNA – which did not differ with treatment in non-fostered males – were significantly increased in D→ fC males (Extended Fig. 8). The importance of these increases remains to be determined, but these findings suggest that cross-fostering impacts the gut epithelium, albeit not identical to the changes that were previously observed.

Finally, we conducted social behavior testing to assess the ability of cross-fostering to a CON dam at birth to rescue social behavior deficits in DEP/MS-exposed male offspring during adolescence. In the sociability assay, D → fD offspring displayed no preference for a social stimulus in the sociability assay (Fig. 5g), similar to DEP/MS offspring that remained with their birth mother (see Fig. 1). However, D→fC males showed a significantly higher preference for a social stimulus (Fig. 6h) and spent significantly more time in social vs. object investigation as compared to D → fD offspring (Fig. 6i), demonstrating that cross-fostering to a CON dam at birth does, indeed, prevent sociability deficits in DEP/MS-exposed males. In the social novelty preference test, D→fC males spent significantly more time in total social investigation as compared to D → fD males, but there was no significant effect on social novelty preference (Extended Fig. 9). This finding may suggest that manipulation of the gut microbiome increases social motivation across assays, rather than choice of social partner (novel vs. cage mate). We also compared the sociability of CON-exposed males fostered to DEP/MS dams (C→ fD) to that of CON-exposed males fostered to CON dams (C→ fC) to test whether exposure to a DEP/MS dam at birth could induce social deficits. We found no difference in any measures of the sociability assay between these groups, suggesting that cross fostering can restore social behavior following DEP/MS exposure but is insufficient to induce a DEP/MS behavioral phenotype on its own (Extended Fig. 9). In sum, these findings demonstrate that intervening at the level of the microbiome can ameliorate social deficits in male offspring following DEP/MS.

### DEP/MS does not alter the maternal gut, vaginal, or milk microbiome

Newborn offspring acquire microbes from their mother during vaginal delivery, as well as from exposure to maternal fecal microbes in the home cage and maternal milk microbes through nursing. We hypothesized that differences in the gut microbiome between CON and DEP/MS male offspring would be due to differential transmission of maternal gut microbes during the perinatal period. To thoroughly characterize the composition of the maternal gut microbiome throughout pregnancy in both treatment groups, we collected fecal boli at baseline (prior to pregnancy) and then every three days throughout gestation in dams from both CON and DEP/MS treatment conditions. We also collected vaginal samples at E17.5 (the eve before birth) as well as milk microbiome samples at P12/13 (peak milk production, Fig. 7a for schematic). To our surprise, we found that while the composition of the gut microbiome shifted significantly over the course of pregnancy, becoming more evenly distributed, there were no differences in the maternal gut microbiome between DEP/MS or CON dams at any timepoint, in either alpha (Fig. 7b) or beta diversity (Fig. 7c). Similarly, there were no differences in the composition of the gut microbiome between DEP/MS and CON foster dams (Fig. 7d&e). Furthermore, we found no differences in the vaginal microbiome (Fig. 7f&g) or milk microbiome (Fig. 7h&I). While counter to our initial hypotheses, these findings point to the intriguing possibility that the signal that leads to microbiome restoration in DEP/MS pups fostered to a CON dam may be conveyed in the non-microbial constituents of milk but is not driven specifically by milk microbes. These data present an exciting avenue for future studies and potential therapeutic development.

**Figure 7.**
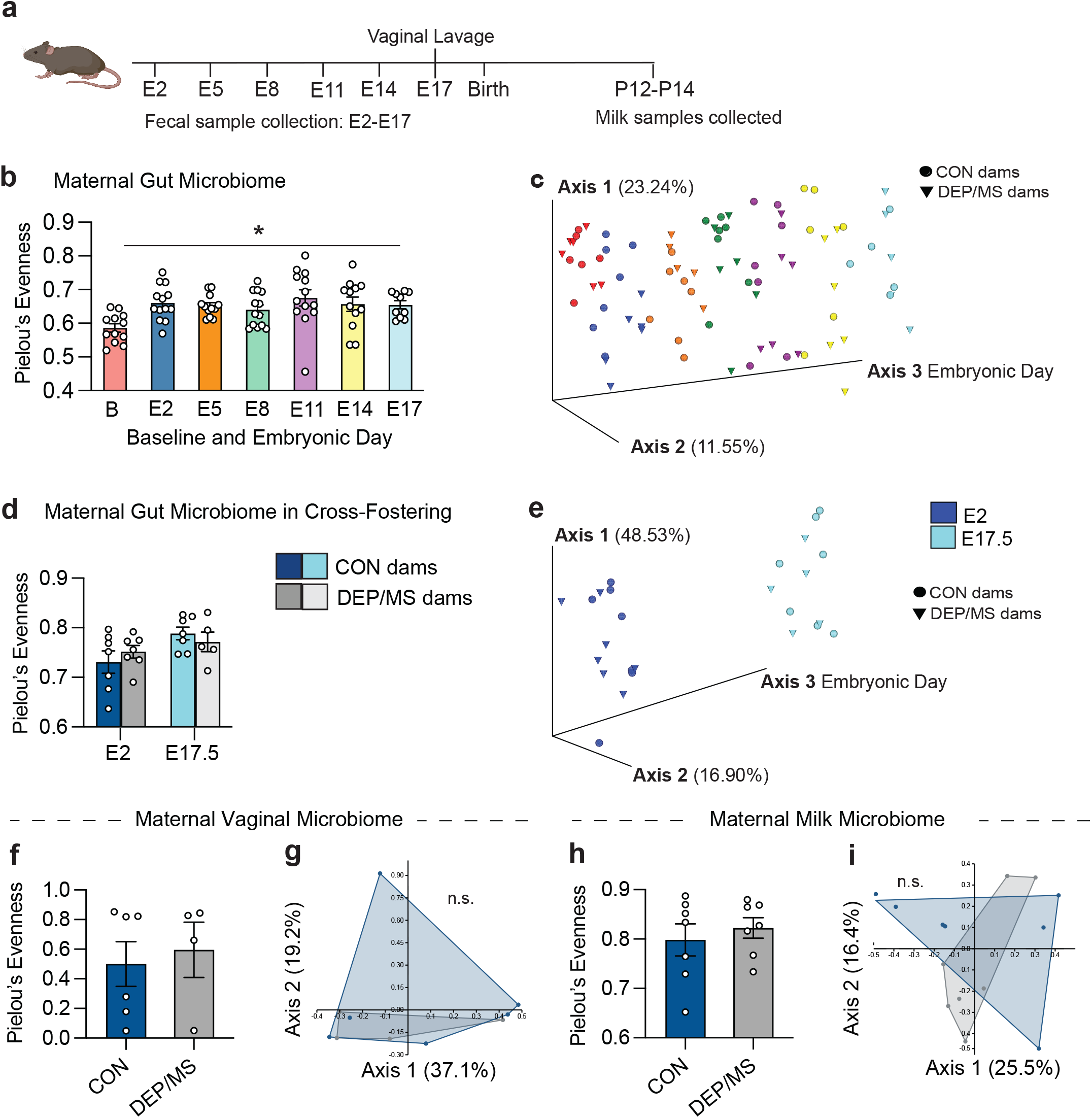
DEP/MS does not impact the maternal gut, vaginal, or milk microbiomes. **a,** Schematic depicting timeline for microbiome sample collections during pregnancy. **b,** In CON and DEP/MS dams, alpha diversity increased significantly over the course of pregnancy (N=5-7/group, Linear Mixed-effect Model first-differences from Pielou’s Evenness, p<0.05), but microbiome evenness did not differ between CON and DEP/MS dams at any timepoint. **c,** Beta diversity trended towards a significant difference with embryonic day (Linear Mixed-effects using first-distances based on weighted UniFrac dissimilarity index, p=0.06), but did not differ between CON and DEP/MS dams. **d**, In the dams used in cross-fostering experiments (CON and DEP/MS), there were no differences in alpha diversity at the beginning (E2) or end (E17.5) of pregnancy (N=5-7/group, Pielou’s evenness, E2: p=0.65, E17.5: p=0.68). **e**, There were also no changes in gut microbiome community structure between CON and DEP/MS dams (N=5-7/group, weighted Unifrac, E2: p=0.68, E17.5: p=0.34). There were no differences between CON and DEP/MS dams in alpha (**f**, N=4-6/group, Pielou’s evenness: p=0.67) or beta diversity (**g,** N=4-6/group, Bray-Curtis: p=0.44) within the vaginal microbiome. Lastly, there were no differences between CON and DEP/MS dams in alpha (**h**, N=7/group, Pielou’s evenness: p=0.75) or beta diversity (**i,** N=7/group, Bray-Curtis: p=0.73) within the milk microbiome. d, f-i data represent Mean +/- SEM, *p<0.05. CON: vehicle/control, DEP/MS: diesel exhaust particles/maternal stress. E: embryonic, P: postnatal.

## Discussion

While human epidemiological work has incontestably shown that environmental toxicants and psychosocial stressors converge on vulnerable populations and that such exposures increase disease burden, little mechanistic work has been done to understand the biological substrates on which these exposures converge. Here, we show that prenatal DEP/MS alters the composition of the gut microbiome, induces microglial hyper-ramification, and decreases dopaminergic tone within the reward circuits of the brain, all in a male-biased manner. Furthermore, we find that social behavior deficits in male offspring following DEP/MS can be rescued or prevented by activating the dopamine system or by shifting the composition of the gut microbiome, respectively.

Our findings point towards both the dopamine system and the gut microbiome as important therapeutic targets in the treatment of social deficits in ASD, as well as other disorders with an environmental component to their etiology. This is in line with previous work in other mouse models which suggests that modification of the gut microbiome can be efficacious for increasing sociability – and that it does so by acting on the dopamine system. Specifically, germ-free mice exhibit robust social deficits which can be rescued by bacterial colonization (Desbonnet et al., 2014). Furthermore, in mouse models of maternal high-fat diet and in genetic models such as Shank 3b KO and Catnap2 KO mice, supplementation of the gut microbiome with specific bacterial species increases social behavior, often by inducing neuroplasticity in the VTA (Buffington et al., 2021; Sgritta et al., 2019). Similarly, symptoms of Parkinson’s Disease (a disorder primarily of dopaminergic neurons) can be induced in mice via colonization with human gut microbes from patients with Parkinson’s (Sampson et al., 2016).

Brain region-specific modification of dopaminergic function remains difficult in human patients. In contrast, the gut microbiome represents a much more tractable therapeutic target in humans. One strength of our work is that our use of a postnatal cross-fostering procedure demonstrates that shifting the overall composition of the gut microbiome is effective at restoring sociability. There are several possible mechanisms by which this might be mediated, including changes in microbial metabolites such as short chain fatty acids (SCFAs) and direct vagal nerve activation (Sherwin et al., 2019). Future studies are needed to determine the precise route by which changes in the gut microbiome following DEP/MS exposure leads to changes in the brain and, therefore, social behavior.

One important point is that while modification of the gut microbiome is sufficient to prevent social behavior impairments in male offspring following DEP/MS, cross-fostering to a DEP/MS dam was insufficient to induce social behavior deficits in CON males. Excitingly, this suggests that the gut microbiome may be an important target for intervention, even when it is not responsible for the entire disease phenotype. Similarly, (Sgritta et al., 2019) found that supplementation with *L. reuteri* rescued social behavior deficits in valproic acid-treated mice, despite *L. reuteri* abundance not being reduced by this exposure in the first place. It also suggests that, at least in part, physiological and behavioral changes in DEP/MS males are organized, at least in part, during gestation. Indeed, the relationship between the host and its bacterial colonizers is bi-directional. One possibility is that changes to the immune system and/or intestinal epithelium in utero precede, and to some degree dictate, which bacterial taxa gain a foothold. This is especially likely given that our effects are observed in male offspring only, despite exposure to the same microbes within the home cage environment. The mechanisms by which this sex-specificity is imparted represent an important area for future investigation.

We observed that cross-fostering to a CON dam on the day of birth prevents social deficits in DEP/MS males, but this does not appear to be driven by differential exposure to maternal microbiome – leaving open the question of what factor is responsible for shifting the microbiome of male offspring. Our previous work has shown that cytokines such as TNFα, IL-17a, and IL-6 are increased in maternal circulation following DEP/MS exposure on E17.5, as is corticosterone (Block et al., 2022). We have also shown that corticosterone is transiently elevated in blood serum of male but not female pups at P1 following DEP/MS exposure (Bolton et al., 2013). One possibility is that differential postnatal exposure to cytokines or hormones in maternal milk might shift the composition of the gut microbiome, and thereby, prevent social behavior deficits. Alternatively, some protective signal may be lacking in DEP/MS milk that is present in the milk of CON dams.

In closing, our results characterize male-specific behavioral, microglial, and neural effects of synergistic exposure to both air pollution and maternal stress during pregnancy. Furthermore, we identify two potential sites for therapeutic intervention in the treatment of social behavior impairments in ASD. In particular, targeting the gut microbiome may be critical to preventing the effects of prenatal environmental toxicant exposures across a number of neuropsychiatric conditions.

## Methods

### Animals

Wild-type (WT) *C57Bl/6J* mice were purchased from Jackson Laboratories (Stock # 000664). DAT-IRES-Cre mice (Jackson Laboratories, Stock # 006660) were obtained from the lab of Dr. Henry Yin at Duke University (used for chemogenetic manipulations). All animals were group housed under standard laboratory conditions (12-hour light/dark cycle [6am-6pm], 23°C, 60% humidity) with same-sex littermates in standard mouse cages. To account for litter effects, male and female offspring were obtained from multiple litters in all treatment groups and all experiments. Complete litter, animal, and number of animals per litter information can be found in Supplementary Table 1. All experiments were conducted in accordance with the NIH *Guide to the Care and Use of Laboratory Animals* and approved by the Massachusetts General Hospital Institutional Animal Care and Use Committee (IACUC), and subsequently, the Duke University IACUC.

### DEP/MS exposures

#### DEP instillations

Diesel exhaust particles (DEP) were obtained from Dr. Ian Gilmour at the Environmental Protection Agency and exposures were conducted in accordance with Bilbo et al. (2013). Multiparous C57Bl/6J mice females were used for all experiments. Timed, harem breeding was performed (2 females housed with 1 male). Females were checked daily for the presence of a vaginal plug, which was taken as an indication of pregnancy and set as embryonic day E0. Females were then pair-housed based on embryonic day until E13. DEP instillations occurred every 3 days throughout pregnancy, beginning on E2, for a total of 6 instillations. Briefly, females were weighed and then anesthetized with 2% isoflurane and administered either 50 μg of DEP/50 μL of vehicle (0.05% Tween20 in PBS; DEP), or vehicle (CON) via oropharyngeal instillation. All females were monitored until they were awake and alert and then returned to their home cages. The concentration of DEP used in this study has been validated and published in several previous manuscripts (Block et al., 2022 BioRxiv, Bolton et al., 2013; 2014; 2017).

#### Maternal Stress (MS)

To induce prenatal maternal stress, we utilized an adapted nest restriction paradigm (Bolton et al., 2013; Rice et al., 2008). On the evening of E13, all females were singly housed. In the MS condition, females were placed into a cage containing a thin layer of alpha-dri bedding (AlphaDri; Shepherd Specialty Papers) covered by an elevated aluminum mesh platform (0.4 cm × 0.9 cm mesh; McNichols Co., Tampa, FL) and given 2/3 of a cotton nestlet (^~^1.9 grams). In the control condition, females were placed into a cage with plenty of Alpha-Dri bedding and provided with an entire cotton nestlet. On E18.5 (evening before pups are born), all dams are placed into a clean cage of Alpha-Dri bedding along with a full cotton nestlet. All offspring were weaned into standard mouse cages with same sex littermates at postnatal day (P)24.

### Behavioral Testing

All behavioral testing took place between postnatal days P27-P40. For all behavioral assays, males and females were tested using separate testing boxes on different days. Animals were handled prior to behavioral testing and habituated to the testing room and testing apparatus one day prior to testing. All behavioral testing was conducted during the second half of the light phase (afternoon).

#### Sociability and Social Novelty Preference

To assess the preference of experimental animals to investigate a social vs a non-social stimulus (Sociability Test), or a novel social vs. a familiar social stimulus (Social Novelty Preference Test) a 3-chambered social preference task was used according to Smith et al., 2015. In this task, the testing apparatus consists of a 3-chambered arena with openings to allow for passage from between the chambers. In the Sociability Test, stimuli (either a novel sex-, treatment-, and age-matched conspecific or a novel rubber duck) were confined within smaller containers composed of Plexiglass rods in each of the opposite side chambers. Subject animals were placed into the middle chamber and their movement and investigation of each of the stimuli was scored over the course of 5 min. In the Social Novelty Preference, the same procedure was followed, except stimuli consisted of either a novel sex-, treatment-, and age-matched conspecific or a familiar cage mate (also matched for sex, treatment, and age) and investigation was quantified over the course of 10 min. The testing apparatus was cleaned with a disinfectant between each test. All videos were scored using Jwatcher (JWatcher.ucla.edu) or Solomon Coder (Solomon.andraspeter.com) by a blinded observer. Scored elements included: time spent in each chamber, time spent investigating each stimulus (i.e. direct sniffing or nose-poking between the bars of the smaller stimulus containers), and time spent in the middle empty chamber. To quantify social preference, ‘sociability’ was calculated as a proportion of investigation time spent investigating the social stimulus ((social investigation time / (social + object investigation time)) x 100). Social novelty preference was calculated as a proportion of investigation time spent investigating the novel social stimulus ((novel investigation time / (novel + familiar investigation time)) x 100). Animals were removed from further analysis prior to unblinding if fighting was noted in the home cage immediately prior to testing (1 instance) or if the wrong stimulus animal was used (1 instance).

#### Marble Burying

As a measure of repetitive behavior, male and female mice were each placed into a clean, standard mouse cage filled with a 5cm depth of wood shavings. 20 blue and black marbles were arranged on top of the bedding in a 4×5 grid. Pictures of each cage were taken prior to introduction of the subject mouse. 20 min. later, each mouse was removed, and a second picture was taken, from which the number of buried marbles was counted by an observer blind to the treatment condition of all animals. A marble was considered “buried” if more than 2/3 of its surface was no longer visible. This test was conducted in a quiet behavioral testing room with dim lighting.

#### Open Field

Open field testing was conducted as a measure of ‘anxiety-like’ behavior. Male and female mice were placed into an open arena (45cm x 45cm). A central square within the arena was outlined on the bottom of the test (15cm x 15cm). Ethovision software (Noldus) was used to quantify total distance traveled, velocity, and time spent in the central square over the course of 10 minutes.

### Microglial isolations

To isolate microglia from the NAc for RNA sequencing, microglial isolations were conducted according to Hanamsagar et al. 2017 and Bordt et al., 2020. Briefly, microglia were isolated from bilateral NAc tissue dissections at P60. Following perfusion with ice cold saline, brains were removed and NAc tissue from both hemispheres was dissected from whole brains on ice using sterile forceps and minced with a razor blade. Homogenate was then submerged in Hank’s Buffered Salt Solution (HBSS; Thermofisher Scientific) containing collagenase A (Roche, 1.5mg/ml) and DNAse 1 (Roche, 0.4 mg/ml). This was then incubated in a water bath at 37°C for 15 min. Next, mechanical digestion of tissue was performed by sequentially passing samples through successively smaller glass Pasteur pipettes. Once a single cell suspension was obtained, samples were filtered, rinsed in HBSS, and centrifuged at 2400 rpm for 10 min at 4°C. Samples were then incubated for 15 min with CD11b antibody conjugated magnetic beads (Miltenyi Biotec) before passage through a magnetic bead column (Quadro MACS Separator and LS columns, Miltenyi Biotec) to separate CD11b+ cells (microglia) from CD11b-cells. Both CD11b+ and CD11b-cell populations were washed in 1X PBS and stored at −80°C until RNA extraction. This is a well-established method for isolating microglia from brain tissue (Bordt et al., 2020).

### Microglial RNA sequencing

Following microglial isolations, RNA from both CD11b+ (microglia) and CD11b-cell populations was extracted according to the Trizol based method described above. Samples were stored at −80°C and then transferred to the MGH Next Generation Sequencing Core for sequencing. Following quality control, NGS libraries were constructed from total RNA using polyA selection followed by NEBNext UltraDirectional II protocol (New England Biolabs) and sequenced on an Illumina HiSeq 2500 instrument, resulting in 20-30 million reads per sample. The STAR aligner (Dobin et al., 2013) was used to map sequencing reads to the mouse mm9 reference genome. Counts for exonic reads were generated using featurecounts (Liao et al., 2014), and non-protein-coding genes were filtered out before any analysis. Data was separated between males and females and Cd11b+ and Cd11b-. Only genes that had 1 count or more in all 4 samples per group were used for differential gene expression analysis using DESeq2 (Love et al., 2014). Custom R scripts and EnhancedVolcano (Blighe et al., 2021) were used to generate the volcano plots. Gene set enrichment analysis (GSEA) was performed using the R package WebGestaltR (version 0.4.4). Enrichment was assessed in the following databases: (1) geneontology_Biological_Process_noRedundant, and (2) geneontology_Cellular_Component_noRedundant. Default settings for GSEA enrichment were employed in the WebGestaltR command except for minNum (0.2) and fdrThr (0.05). Analysis was performed on a ranked list of differential genes that was determined by the equation sign(log2FoldChange)*-log10(p-value) from the DESeq2 output. Positive rank scores reflected genes up-regulated in the DEP/MS microglia, while negative rank scores indicated genes up-regulated in control microglia. Pathways shown in figures are Biological Processes (FDR<0.05), and Cellular Components (FDR<0.05).

Stratified ranked-ranked hypergeometric overlap (RHHO) analysis was done using the R package RRHO2 (version 1.0), and default settings in the RRHO2_initialize command. To allow for comparison with published data which often reported normalized gene counts, gene rank values were calculated by first performing unpaired t-tests between normalized gene expression values between experimental conditions. The average values of the experimental groups were then used to calculate a fold-change in mean expression. To obtain the gene rank, the sign of this fold-change was multiplied by the log10(p-value). Published datasets used for this analysis are cited above in the results section. All code for processing is currently hosted and publicly available (github/XXXXXX). All data will be publicly available; GEO Accession number pending.

### Immunohistochemistry (IHC) staining and analysis

For all IHC, animals were euthanized via CO_2_ inhalation and brains were perfused with ice-cold saline followed by 4% paraformaldehyde. Brains were then removed and post-fixed for 48 hrs in 4% paraformaldehyde, followed by 48 hrs in 30% sucrose with 0.1% sodium azide. Subsequently, brains were flash frozen in 2-methylbutane and stored at −20°C until sectioning at 40μm on a cryostat (Leica Biosystems) into cryoprotectant.

#### Ionized calcium-binding adaptor molecule (Iba1) for microglial density/morphology

Free-floating sections containing NAc were rinsed 3x in PBS, incubated in 10 mM sodium citrate for 30 minutes at 75 ºC for antigen retrieval, and rinsed 3x in PBS again. They were then blocked for 1hr in PBS with 1% H_2_O_2_, 10% normal goat serum, and 0.3% Triton-X. Next, sections were incubated with primary antibody (rabbit anti-Iba1, 1:5000; Wako Chemicals) overnight at RT on a shaker. Following 3x PBS rinses, sections were then incubated in biotinylated secondary antibody (goat anti-rabbit IgG, 1:500; Vector Laboratories) for 1hr at RT. Immunostaining was then amplified and identified using the streptavidin/HRP technique (Vectastain ABC kit; Vector Laboratories) with diaminobenzidine (DAB; Vector Laboratories) as the chromagen. Sections were mounted on gelatinized slides, dehydrated, and coverslipped with Permount (Fisher Scientific).

Digitized images of 20X tissue section were analyzed using FIJI software (https://imagej.net/software/fiji/). Brightfield sections were imaged using a Zeiss microscope with apotome attachment (Axio Imager Z1). Two images per hemisphere per animal, in the portion of the NAc directly medial to the anterior commissure (bregma 1.33 mm), were used for mean pixel intensity analysis. Automatic thresholding and mean pixel intensity were performed using ImageJ software. For 3D Morph analysis of microglial morphology, two brightfield z-stacks (10 images per stack) per section within the NAc (directly medial to the anterior commissure; Bregma 1.33mm) were acquired on a Zeiss microscope with apotome attachment (Axio Imager Z1) with 40X magnification and a 0.65 μm step size. Thresholding was performed on each z-stack using FIJI, and the background was inverted for each image. These files were then uploaded into MATLAB. Microglial morphological analysis was performed using 3DMorph, a MATLAB-based script that semi-automatically processes individual microglial morphology from overlapping 3D clusters (York et al., 2018). 3DMorph open-source code was provided and downloaded from GitHub (https://github.com/ElisaYork/3DMorph).

#### Dopamine D1 receptor (D1R) and tyrosine hydroxylase (Th) in the NAC at P45

Separate sections were used for D1R and Th. Free-floating sections containing NAc were rinsed 3x in PBS, incubated in 10 mM sodium citrate for 30 minutes at 75 °C for antigen retrieval, and rinsed 3x in PBS again. For D1R staining only, sections were also incubated for 1hr in 50% methanol followed by 1hr in Sodium Tetraborate (1mg/ml in 0.1M PB). All sections (D1R and Th) were then blocked for 1hr in PBS with 1% H_2_O_2_, 10% normal goat serum, and 0.3% Triton-X. Next, sections were incubated with primary antibody (D1R: anti-D1R [mouse] 1:1500, Novus Biologicals, 48 hrs at 4°C; Th: anti-Th [rabbit] 1:1000, ThermoFisher Scientific, overnight at RT on a shaker). Following 3x PBS rinses, sections were then incubated in secondary antibody (D1R: goat anti-mouse Alexa-Fluor 488, 1:500, Thermofisher Scientific, 4 hrs at RT; Th: goat anti-rabbit Alexa-Flour 488, 1:500, Thermofisher Scientific, 4 hrs at RT). Sections were mounted on gelatin subbed slides and coverslipped with VectaShield anti-fade mounting medium (Vector Labs).

For analysis of each (D1R and Th), a Zeiss AxioImager microscope was used to take 40x magnification Z-stacks in the NAc. 10-step Z-stacks were taken with a step-size of 0.65μm measured from the center of the image. Images were taken medial to the anterior commissure (AC) using the AC as a landmark. A total of 3-6 images were taken for each animal. Mean grey value of fluorescent intensity was analyzed using FIJI. Briefly, Z-stacks were converted to maximum projection images and mean grey value was measured and normalized to background (defined as the mean grey value of the AC). The values of all images taken for a given animal were averaged to provide a single measurement per animal.

#### D1R and Iba1 co-labeling at P30 in males only

Sections containing the NAc were rinsed 3x in PBS, incubated in 10 mM sodium citrate for 30 minutes at 75 ºC for antigen retrieval, and rinsed 3x in PBS again. Sections were then blocked for 1hr in PBS with 1% H_2_O_2_, 10% normal goat serum, and 0.3% Triton-X followed by 1hr in 50% methanol and 1hr in Sodium Tetraborate (1mg/ml in 0.1M PB). Next, sections were incubated with primary antibody (D1R: anti-D1R [mouse] 1:1500, Novus Biologicals; Iba1: anti-iba1 [chicken], 1:1500 Synaptic Systems) for 48hrs at 4°C. Following 3x PBS rinses, sections were then incubated in secondary antibody (D1R: goat anti-mouse Alexa-Fluor 488, 1:500, Thermofisher Scientific; Iba1: goat anti-chicken Alexa-Fluor 568, 1:500, Thermofisher Scientific) for 4 hours at RT. Sections were mounted on gelatin subbed slides and coverslipped with VectaShield anti-fade mounting medium (Vector Labs).

To quantify microglial engulfment of D1R, 63X magnification Z-stacks were taken on a Ziess Airyscan 880 Confocal Laser scanning microscope. Z-stacks ranged in size depending on the size of the microglia, but with a consistent step size of 0.3μm. Imaris 9.5.1 (Bitplane Scientific Software) was used to create surface renderings of individual microglia (Iba1 labeling) and D1R (D1R labeling) within the microglial surface. Volume of engulfed D1R was then quantified and normalized to total cell volume. IMARIS filament tracer was used to assess morphological characteristics of microglia in 3D and to generate Sholl analyses. 4-5 cells were reconstructed per animal from 3-5 separate brain sections within the NAc.

#### Th and mCherry co-labeling in the ventral tegmental area (VTA; for verification of viral transfection)

Free-floating sections containing VTA were rinsed 3x in PBS, incubated in 10 mM sodium citrate for 30 minutes at 75 °C for antigen retrieval, and rinsed 3x in PBS again. Sections were then blocked for 1hr in PBS with 1% H_2_O_2_, 10% normal goat serum, and 0.3% Triton-X. Next, sections were incubated with primary antibody (anti-Th [mouse] 1:1500, Immunostar; anti-dsRed [rabbit] 1:1500, Clontech) overnight at RT shaking. Following 3x PBS rinses, sections were then incubated in secondary antibody (Th: goat anti-mouse Alexa-Fluor 488, 1:500, Thermofisher Scientific, dsRed: goat anti-rabbit Alexa-Flour 568, 1:500, Thermofisher Scientific) for 4 hours at RT. Sections were mounted on gelatin subbed slides and coverslipped with VectaShield anti-fade mounting medium (Vector Labs). 10X images were taken on a Zeiss AxioImager microscope to co-labeling of Th and dsRed.

### Tissue Punches for Neural Gene Expression

#### Tissue Collection and Punches

To quantify gene expression for socially relevant receptors in discrete brain regions, tissue punches followed by qPCR were used. At P45, animals were euthanized via C02 inhalation and transcardially perfused with ice cold saline. Brains were removed and snap frozen in 2-methylbutane. Punches from the NAc and amygdala were collected using a 1mm diameter core sampling tool (Electron Microscopy Sciences). Brains were mounted in a sterilized cryostat and punches were collected by inserting the sterilized core sampling tool to a depth of 1mm. Sections from brains were collected onto slides to verify location and depth of punches. Punches were immediately placed into Trizol on dry ice and transferred to −80C freezer until RNA extraction.

#### RNA Extraction, cDNA synthesis, and qPCR

Samples were homogenized in Trizol® (Thermo-Fisher Scientific) and then vortexed for 10 min at 2000 rpm. After 15 min resting at room temperature (RT), chloroform was added (1:5 with Trizol) and samples were vortexed for 2 min at 2000 rpm, allowed to sit at RT for 3 min, and then centrifuged (15 min at 11,800 rpm; 4°C). From the resulting gradient, the aqueous phase was separated, and isopropanol was added to precipitate RNA (1:1 with aqueous phase). Samples were again vortexed, allowed to set at RT for 10 min, and then centrifuged (15 min at 11,800 rpm; 4°C). Pellets obtained after this step were rinsed twice in ice-cold 75% Ethanol and then resuspended in 8 μl of nuclease-free water. RNA was frozen at −80°C until cDNA synthesis. cDNA was synthesized using the QuantiTect Reverse Transcription Kit (Quiagen). Briefly, RNA (200ng/12 μl nuclease-free H_2_O) was pre-treated with gDNase at 42°C for 2 min to remove genomic DNA contamination. Next, master mix containing both primer-mix and reverse transcriptase was added to each sample and all samples were heated to 42°C for 30 min and then 95°C for 3 min (to inactivate the reaction) in the thermocycler. qPCR was run on a Mastercycler ep realplex (Eppendorf) using the SYBR Green PCR Kit (Quiagen). All PCR primers were designed in the lab and purchased from Integrated DNA technologies. Relative gene expression was calculated using the 2-ΔΔCT method, relative to the house-keeping gene (*18S*) and the lowest sample on the plate (Williamson et al., 2011; Livak & Schmittgen, 2001). Samples were removed before unblinding if they failed to amplify or if there was a secondary peak in the melting temperature plot indicating contamination.

Primer sequences for all genes are as follows: ***18S**, F: GAA TAA TGG AAT AGG ACC GC, R: CTT TCG CTC TGG TCC GTC TT; **Drd1**, F: GAG TGA TTG GGG GAA GTC TG, R: GAC AGG ATA AGC AGG GAC AG; **Drd2**, F: CCT CCA TCG TCT CGT TCT AC, R: GAG TGG TGT CTT CAG GTT GG; **Oprm1**, F: CAT CCA GAC CCT CGC TAA AC, R: GCC AGA GAG GAA TGA CTT TGA; **Oprk1**, F: GCA CCA AAG TCA GGG AAG AT, R: GGC AAA GAC GAA GAC ACA GA; **Oxtr**, F: AGC AGA CAC ACA CAC CTA TG, R: CAG AAC TCG GCT CTT GAA CT; **Ocln**, F: CTG ACT ATG CGG AAA GAG TTG AC, R: CCA GAG GTG TTG ACT TAT AGA AAG AC; **Zo1**, F: AAG AAA AAG ATG CAC AGA GTT GTT, R: GAA ATC GTG CTG ATG TGC CA; **TNFα**, F: GTC GTA GCA AAC CAC CAA, R: AGA ACC TGG GAG TAG ACA AGG; **IL-1β**, F: GTC TTC CTA AAG TAT GGG CTG, R: CAC AGG CTC TCT TTG AAC; **TLR4**, F: CAG CAG AGG AGA AAG CAT, R: CAC CAG GAA TAA AGT CTC TG; **S100β**, F: CTG TCT ACA CTC CTG TTA CTC G, R: CTG CTC CTT GAT TTC CTC CA*.

### Chemogenetic manipulations

DAT-Ires-Cre mice (B6.SJL-Slc6a3^tm1.1(cre)Bkmn^/J, Stock No. 006660, Jackson Laboratory) were obtained from the lab of Dr. Henry Yin. Homozygous males were bred with WT *C57Bl/6* females which were then exposed to DEP/MS during pregnancy according to the methods described above. All resulting offspring were heterozygous (Cre+) and were ear tagged and tail clipped for genotyping at P14 and weaned at P22. On P23 or P24, CON and DEP/MS male offspring underwent stereotaxic surgery to virally inject either pAAV-hSyn-DIO-hM3D(Gq)-mCherry (Excitatory DREADD; Addgene #44361, titers: 7×10^12^ vg/ml) or pAAV-hSyn-DIO-mCherry (Control mCherry reporter; Addgene #50459, titers: 7×10^12^ vg/ml) into the VTA. Coordinates were, AP: - 2.6, ML +/-1.2, DV – 4.8, 10°angle. Coordinates were based on the Franklin & Paxinos Mouse Brain Atlas (Franklin & Paxinos, 6^th^ Edition) and adapted for use in juveniles. In brief, surgical procedures were as follows. Juvenile mice were anesthetized with 2% isoflurane anesthesia and pretreated with a subcutaneous injection of Ketophen anesthetic (5mg/kg). Animals were fitted into a Kopf bilateral stereotaxic apparatus (Kopf Instruments) and, following skin preparation with betadine and shaving, an incision was made into the skin. The surface of the skull was cleaned and Bregma was identified. Distance from bregma to the VTA was measured bilaterally and 1μl Hamilton syringes (Hamilton Company) were lowered to the appropriate depth. Virus was injected in a volume of 0.5μl over the course of 5 min and allowed to sit for 5 min before slowly removing the Hamilton syringe. Following injections, the skin incision was closed with wound clips (EZ clip wound closures, Stoelting Co.) and topped with Neosporin and topical analgesic (Lidocaine). Animals were placed on a heating pad and observed until they regained sternal recumbency. They were then weighed daily for the next 5 days and observed for signs of lethargy, pain, or distress.

On P33-35, after allowing 1 week for recovery and viral transfection to occur, subjects were tested on the sociability assay. Methods were in accordance with those described above. Subjects were injected with either Vehicle (1% DMSO in sterile saline) or CNO (1mg/kg in 1% DMSO in sterile saline) 30 min. prior to behavioral testing. After behavioral testing, all animals were euthanized, and brains were collected to verify transfection of the VTA. Behavior was scored according to the methods described above using Solomon Coder by a blinded observer.

### 16S Microbiome sequencing

Bacterial taxa were identified using 16S rRNA amplicon sequencing of microbiome samples. Library preparation was conducted in accordance with standard protocols (earthmicrobiome.org). First, DNA was extracted from all samples using a DNeasy Powersoil Kit (Qiagen, Germantown, MD). Next, PCR with individually barcoded primers (515F-806R; Parada et al., 2016; Caporaso et al., 2011, 2012) was used to amplify the V4 hypervariable region of the 16S rRNA gene. PCR product was then purified (PCR Purification Kit, Qiagen, Germantown, MD), DNA concentration was measured using a Quant-iT Picogreen Assay (Thermofisher Scientific), and an equimolar pool of all samples was made and transferred to the core (MGH: MGH Next-Generation Sequencing Core, Duke: Duke University Center for Genomic and Computational Biology) for sequencing on an Illumina MiSeq Instrument resulting in 20,000 paired-end 250bp reads per sample (Illumina, San Diego, CA, USA).

Qiime2-2020.2 analysis platform, PAST (PAleonto-logical Statistics; Hammer et al., 2001), and the R environment (version 3.4.2) were used to analyze 16S data. Briefly, forward and reverse reads were imported, demultiplexed, and quality filtered using DADA2 (Callahan et al., 2016). Amplicon sequence variants were then aligned with MAFFT (Katoh, 2002) and a phylogenetic tree was generated. PERMANOVA tests with 999 permutations were used to test for between sample differences in bacterial community composition, using unweighted and weighted UniFrac, Bray Curtis, and Jaccard dissimilarity indices. Differences in alpha diversity measures of richness and evenness were tested by Kruskal-Wallis. Longitudinal analysis of beta and alpha diversity indices was conducted using linear mixed effects modeling (LME) with a random-intercept by subject on first-distances and first-differences (Bokulich et al., 2018). Taxonomy was assigned using a Naïve Bayes filtered classifier trained on the Greengenes database, version 13_8, at 99% sequence similarity (McDonald et al., 2012). Differences in relative abundance were assessed using ANCOM and *p* values were adjusted to control for a false discovery rate of 5% to account for multiple comparisons. The extent of differences in the taxonomic profiles between treatment conditions was substantiated further by analyzing linear discriminant analysis of effect sizes calculated through the algorithm from LEfSe (Segata et al., 2011).

### Milking Procedure

Mouse dams were exposed to CON or DEP/MS as described above. Dams were milked on P12, P13, and P14 when milk production is highest. Pups from dams that were milked were not used for any analyses. Milking always took place during the first half of the light phase (9am-12pm). 1 hr prior to milking, dams were separated from their pups within their home cage by placing the pups in a modified igloo (entrances taped so that dams could not enter). Dams were then anesthetized with 2% isoflurane and injected subcutaneously with oxytocin (2IU/kg) to stimulate milk expression. The hair around the nipples was trimmed with fine scissors. Milk was collected into 1.5ml Eppendorf tubes by manually expressing drops of milk into microhematocrit capillary tubes (Fisher Scientific). Milk was immediately placed on ice, spun briefly, and stored at −80°C until further analysis.

### Gut tissue collection, gene expression, and staining

For all gut tissue collection, mice were euthanized via CO_2_ inhalation and transcardially perfused with ice-cold saline. Intestines were removed and placed into a petri-dish of sterile 1X PBS. ^~^1.5cm segments from the duodenum, ileum, and colon were collected. Duodenum was collected proximal to the stomach, and ileum and colon were collected proximal to the cecum. Gut sections were gently compressed to clear them of contents, and either: a) flash frozen in liquid nitrogen for mRNA analysis, b) postfixed for 48 hours in 4% paraformaldehyde followed by 30% sucrose for IHC, or c) postfixed for 5 days in 4% paraformaldehyde followed by 70% Ethanol for H&E staining. After collection, mRNA analysis was conducted according to the methods described above.

For hematoxylin and eosin (H&E staining), gut sections were embedded in paraffin and sectioned at 20μm onto slides. To conduct the staining, slides were first deparaffinized in xylene, rinsed 2x in 100% ethanol followed by 2x in 95% ethanol (5 min. each), and then rinsed in H_2_O. Next, slides were placed in Harris hematoxylin for 15 min., dipped once in 70% acidic alcohol, dipped once in 1% lithium carbonate, and then rinsed in warm H_2_O. Finally, slides were placed in alcohol Eosin Y for 5 min. before xylene dehydration and coverslipping with cytoseal.

Next, three 20 μm ileum cross sections that were localized 100 μm from each other were evaluated for each animal. Representative images of the H&E histology are taken using an Optronics Magnafire digital camera (Optronics) at an image size of 1388×1040 pixels on a Zeiss Axioskop microscope (Carl Zeiss). For each section assessed, a 5x image is taken to capture an overall view of the gut section for orientation followed by 10x images that capture portions of each gut cross section. From each of these sections morphology was evaluated by identifying and isolating intestinal villi based on the following criteria: a) full cross section of villus was present with visible lamina propria, b) epithelial cell lining of villus was continuous and well-organized, c) there was continuity between the lamina propria space and peri crypt space at the base of the villus without epithelial cells disrupting this continuity, and d) there were no major artifacts obscuring the view of the villus cross section. For each villi identified, the following measurements were taken: 1. Villus length - the distance from the base of the villus above the crypt to the end of the villus tip protruding into the luminal space, 2. Villus Width - the average thickness of the villus, 3. Crypt Length/Depth-The distance from the base of the crypt bordering the muscular layer to the approximate top portion of the crypt, 4. Mucosal Thickness -the distance from the base of the crypt bordering the musucular layer to the villus tip protruding into the luminal space; the combined distance of crypt length and villus length, and 5. Villus/Crypt Ratio - A ratio of villus length in comparison to crypt length. The average of each measurement for each animal was taken.

### Cross-fostering experiments

For cross-fostering experiments, WT C57Bl/6 dams were mated and exposed to CON or DEP/MS according to the methods described above. On the morning of the day of birth (P0), pups were cross-fostered to either a mother of the same treatment (CON-CON [C→ fC] or DEP/MS-DEP/MS [D→ fD]), or to a mother of the opposite treatment (DEP/MS-CON [D→ fC] or CON-DEP/MS [C→ fD]). Entire litters were swapped by removing each litter and placing them in a small paper cup filled with bedding from the recipient mother. The entire litter was then placed into the foster cage at once. Maternal care was assessed 3x daily for the first 3 days of life. Briefly, 10 observations were made of each dam during a 1hr period in the morning, afternoon, and evening (during the dark phase) for a total of 30 observations. The behavior of the dam was recorded at each observation. Behaviors were coded as off nest, off nest eating/drinking, on nest, on nest nursing, on nest licking and grooming. Offspring were weaned at P24 into cages with same sex littermates. Social behavior of male offspring was tested between P30-P35, in accordance with the methods described above. Following behavioral testing, all offspring were euthanized and cecal contents collected for microbiome sequencing. Gut segments were also collected for gut gene expression and H&E staining, according to the aforementioned methods.

### Statistics and Reproducibility

Statistical analyses were conducted using GraphPad Prism 9 software, except in the case of the 16S sequencing analyses which were conducted using the Qiime2-2020.2 analysis platform, PAST (PAleonto-logical STatistics), and the R environment and the microglial RNA sequencing data which was analyzed using the R environment and Python. Detailed descriptions of all statistical methods are available in the figure legends and in Supplementary Table 2. All analysis were 2-tailed. Investigators were blind to treatment group for all data analyses. Animals were excluded from further analyses if they were statistical outliers. No statistical methods were used to predetermine samples sizes, but sample sizes are consistent with our previous reporting of similar endpoints (Block et al., 2022; Kopec et al., 2018). Data are expressed as mean +/- SEM and statistical significance was set at p<0.05.

## Acknowledgements

We would like to thank all the members of the Bilbo lab, past and present, for their helpful discussions and feedback. We would also like to thank the MGH Next-Generation Sequencing Core and the Duke Center for Genomic and Computational Biology for their assistance with RNA and 16S sequencing. We thank the animal care staff at MGH and Duke University for their excellent animal care. We have no conflicts of interest to disclose. This work was supported by R01 ES025549 to SDB, R01 ES033056 to SDB, F32 ES029912 to CJS, K99 ES033278 to CJS, and by the Robert and Donna Landreth Family Foundation.

---

**Extended Figure 1.**
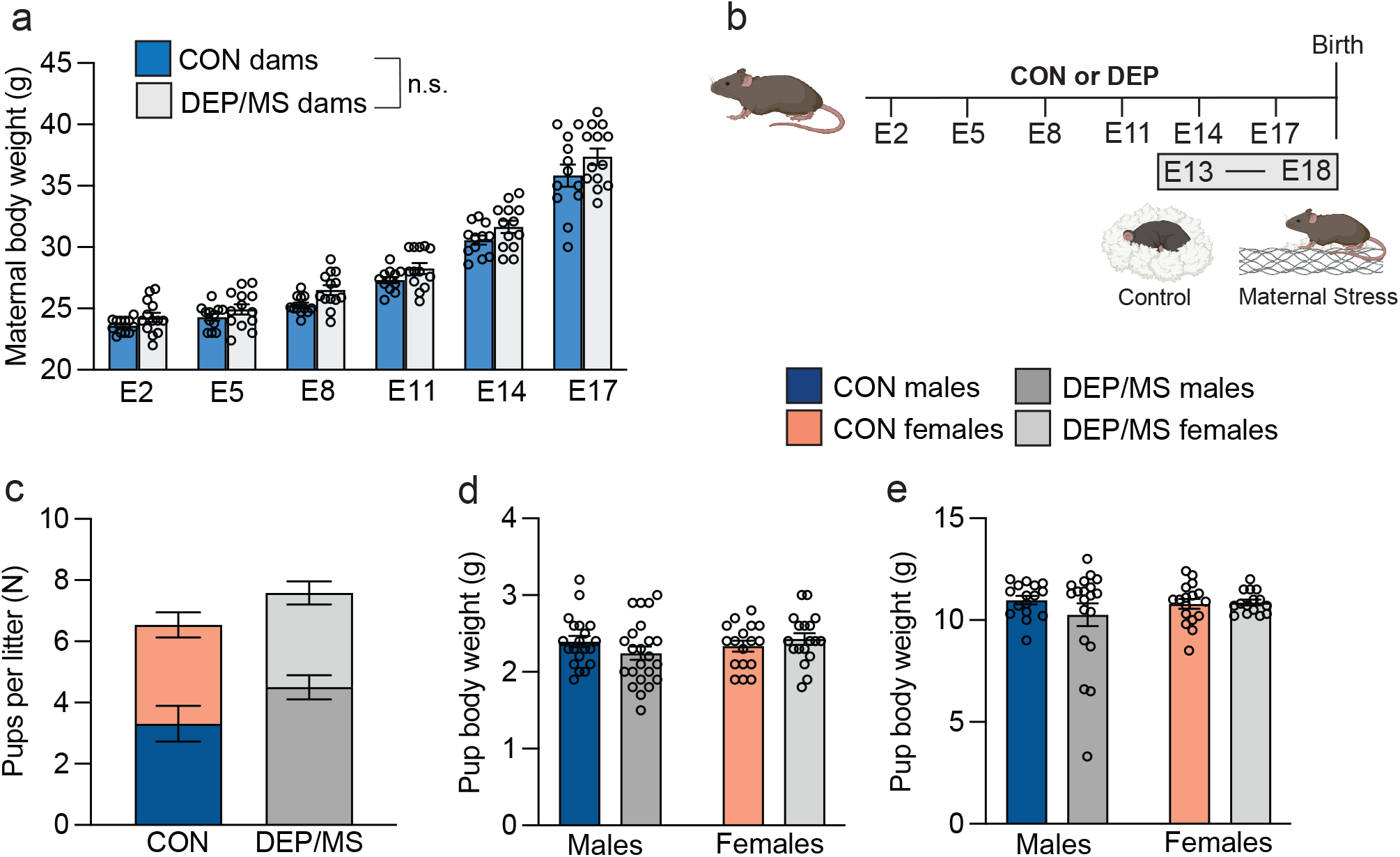
Neonatal outcomes following DEP/MS. **a**, Maternal weight gain across pregnancy. While all dams gained weight across pregnancy, CON and DEP/MS dams do not differ in gestational weight gain (N=12-13/group, 2-way ANOVA [time x treatment], time: p=0.0001, treatment: p=0.06). **b**, Schematic of DEP/MS or CON procedure. **c**, Number of pups/sex/litter does not differ between CON and DEP/MS (N=12-13/group, 2-way ANOVA [treatment x sex], treatment: p=0.26, sex: p=0.69). **d**, There was no significant effect of treatment or sex on body weight at P4 (N=17-23/group, 2-way ANOVA [treatment x sex], treatment: p=0.71, sex: p=0.61) or P24 (N=14-17/group, 2-way ANOVA [treatment x sex], treatment: p=0.80, sex: p=0.57; **e**). Data represent mean + SEM, *p>0.05. CON: control, DEP: diesel exhaust particles, E: Embryonic day.

**Extended Figure 2.**
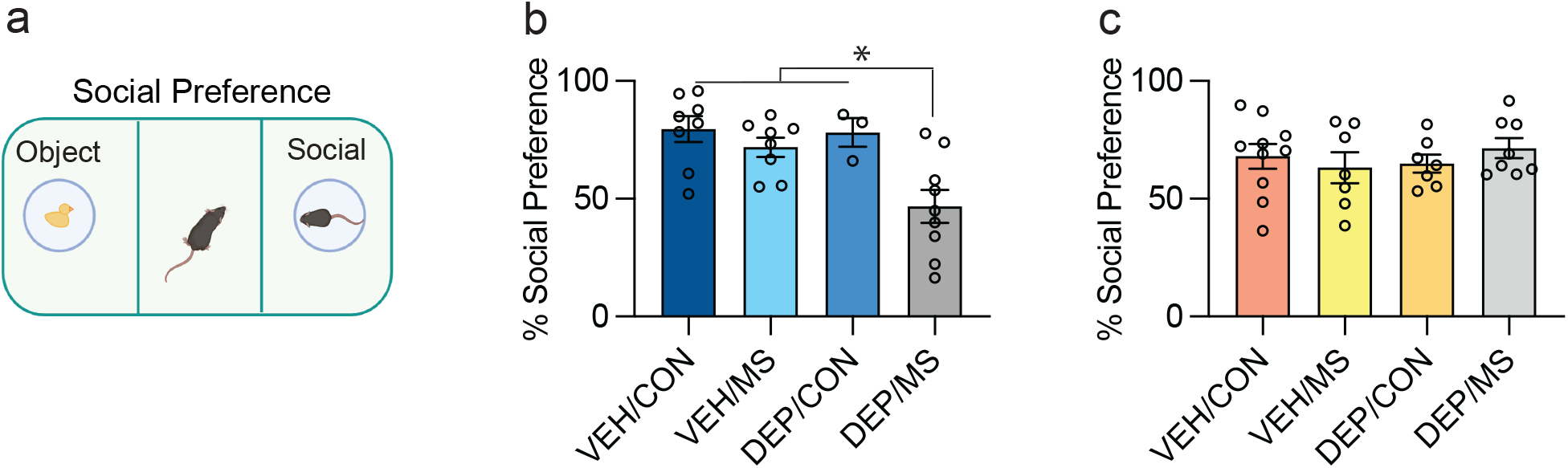
Social behavior following either MS or DEP treatment alone. **a**, Schematic of sociability assay. **b**, In male offspring, neither DEP or MS alone impairs sociability. However, combined DEP+MS exposures eliminates the preference for a social partner (N=3-10/group; one-way ANOVA [treatment], treatment p=0.002) **c**, Neither DEP alone, MS alone, nor DEP/MS alters sociability in females (N=7-10/group; one-way ANOVA [treatment], treatment p=0.71). Data represent mean + SEM, *p<0.05. CON: control, DEP: diesel exhaust particles, MS: maternal stress, VEH: vehicle.

**Extended Figure 3.**
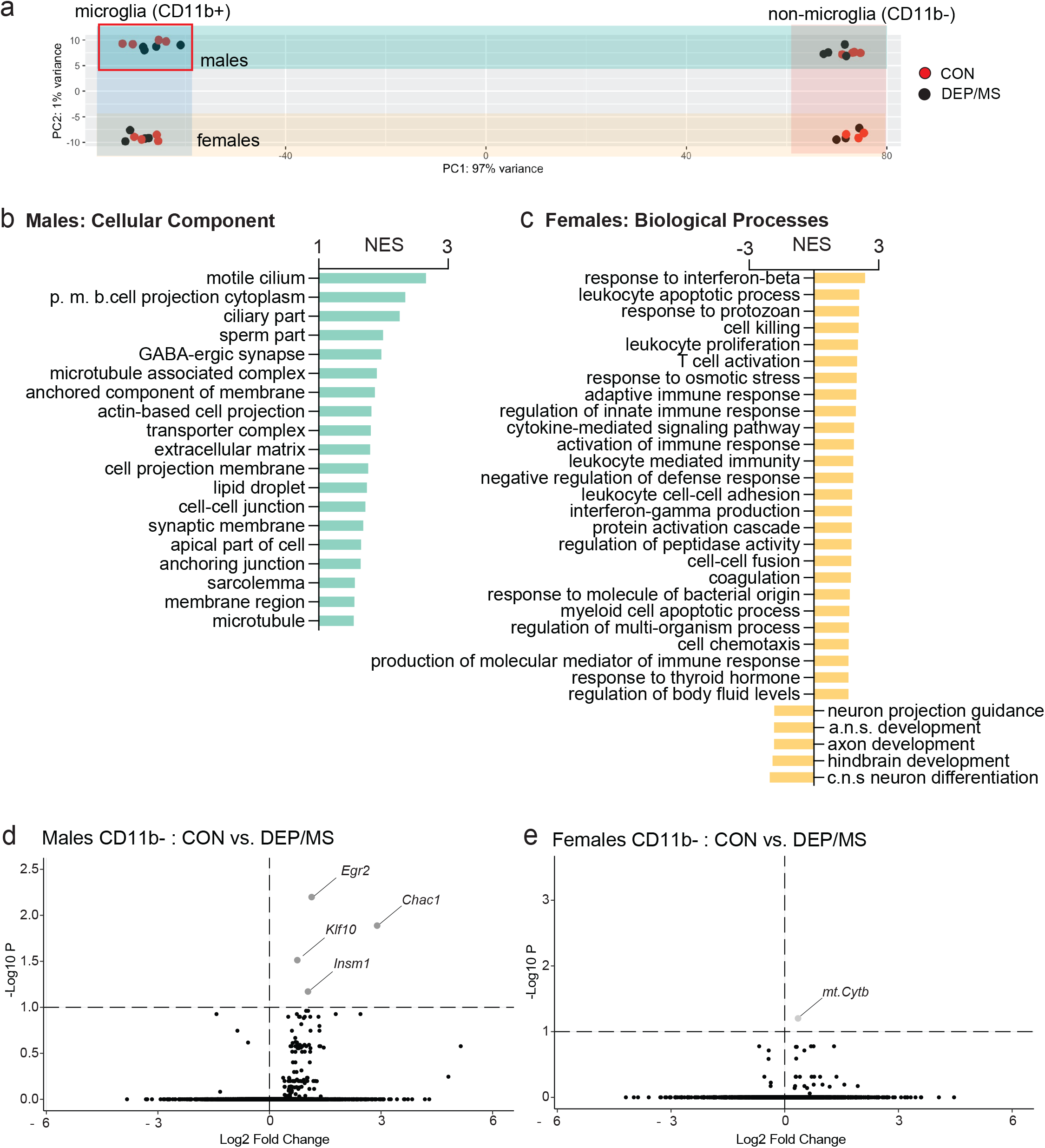
Extended microglial RNA sequencing results. **a**, PCA plots showing distinct clustering of CD11b- and CD11b+ cells, as well as male and female samples within each sample (N=4/group). In male CD11b+ cells, CON and DEP/MS samples are non-overlapping. **b**, GSEA analysis of CD11b+ samples demonstrates changes in cellular components related to cell projection assembly and motility in males following DEP/MS (N=4/group, all pathways shown reached significance with FDR p<0.05). Positive scores indicate up-regulation in DEP/MS relative to CON. **c**, GSEA analysis of CD11b+ samples demonstrates changes in biological pathways related to interferon-ß, innate immunity, and T-cell activation in females following DEP/MS (N=4/group, all pathways shown reached significance with FDR p<0.05). Positive scores indicate up-regulation in DEP/MS relative to CON. **d-e**, Volcano plots of CD11b-samples demonstrate very few genes that differentially expressed in following DEP/MS in males (**d**) and females (**e**, N=4/group, positive Log2 Fold Change indicates higher expression following DEP/MS). CON:vehicle/control, DEP/MS: diesel exhaust particles/maternal stress, N: number, NES: normalized enrichment score, p.m.b.: plasma membrane bound. c.n.s.: central nervous system, a.n.s. autonomic nervous system, PCA: principal component analysis. GSEA: gene set enrichment analysis.

**Extended Figure 4.**
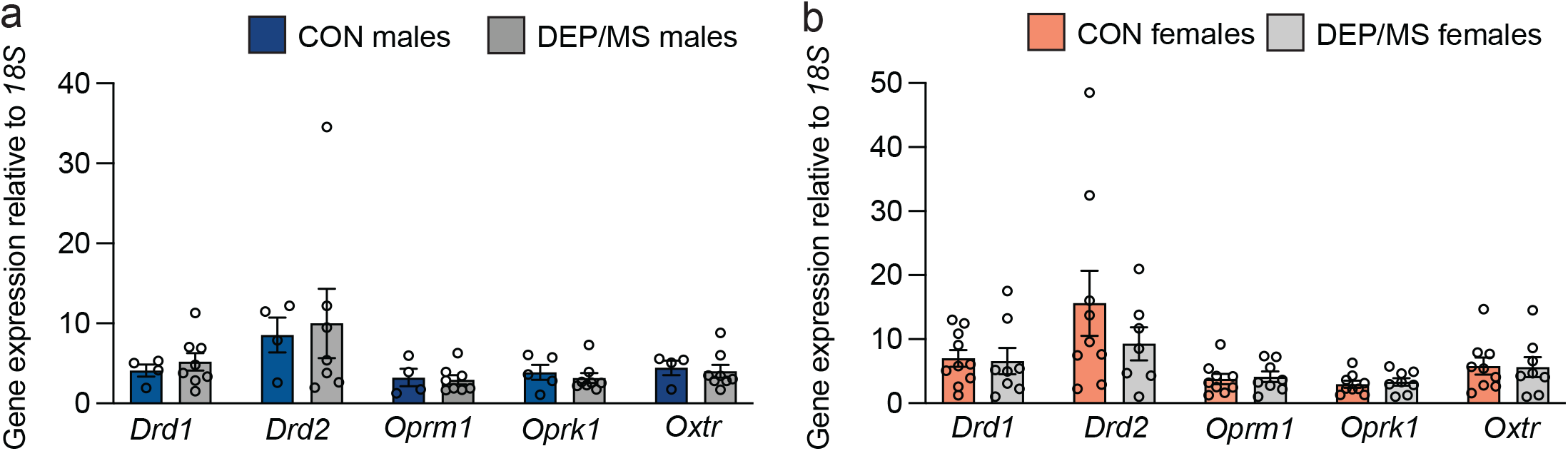
DEP/MS has no effect on gene expression for socially-relevent receptors in tissue punches from the amygdala in either male (**a**, N=4-8/group, unpaired t-tests [CON vs DEP/MS], *Drd1*: p=0.53, *Drd2*: p=0.82, *Oprm1*: p=0.81, *Oprk1*: p=0.53, *Oxtr*: p=0.75) or female (**b,** N=7-10/group, unpaired t-tests [CON vs DEP/MS], *Drd1*: p=0.86, *Drd2*: p=0.33, *Oprm1*: p=0.77, *Oprk1*: p=0.66, *Oxtr*: p=0.92) offspring at PND45. Data represent mean +/- SEM, *p=0.05. CON: vehicle/control, DEP/MS: diesel exhaust particles/maternal stress.

**Extended Figure 5.**
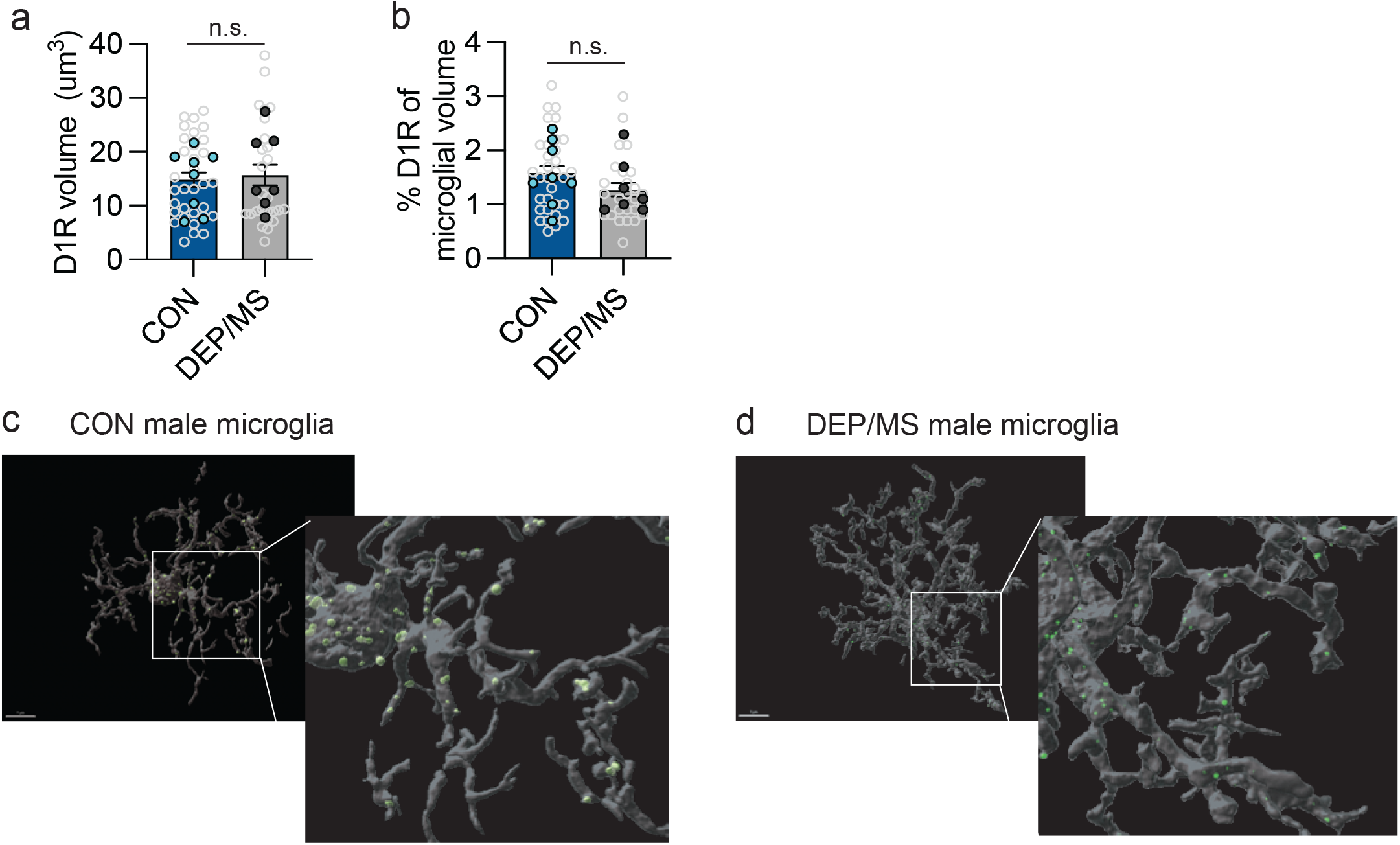
Microglia do not engulf more D1R in the NAc following DEP/MS exposure in males at P30. **a-b,** DEP/MS has no effect on D1R engulfment (as assessed by D1R volume (**a**) and % D1R volume within microglial volume (**b**) within the nucleus accumbens (NAc, N=7-8 animals/group), nested t-tests [microglia per animal; CON vs. DEP/MS], D1R volume: p=0.69, % D1R of microglial volume: p=0.377). Light grey dots represent individual microglia, but data are nested within animals (animal averages in darker dots). **c-d**, Representative Imaris 3D reconstruction of CON (**c**) and DEP/MS (**d**) microglia within the NAc of males at P30, scale=7μms. Data represent mean +/- SEM, *p<0.05. CON:vehicle/control, DEP/MS: diesel exhaust particles/maternal stress, D1R: dopamine D1 receptor, NAc: nucleus accumbens.

**Extended Figure 6.**
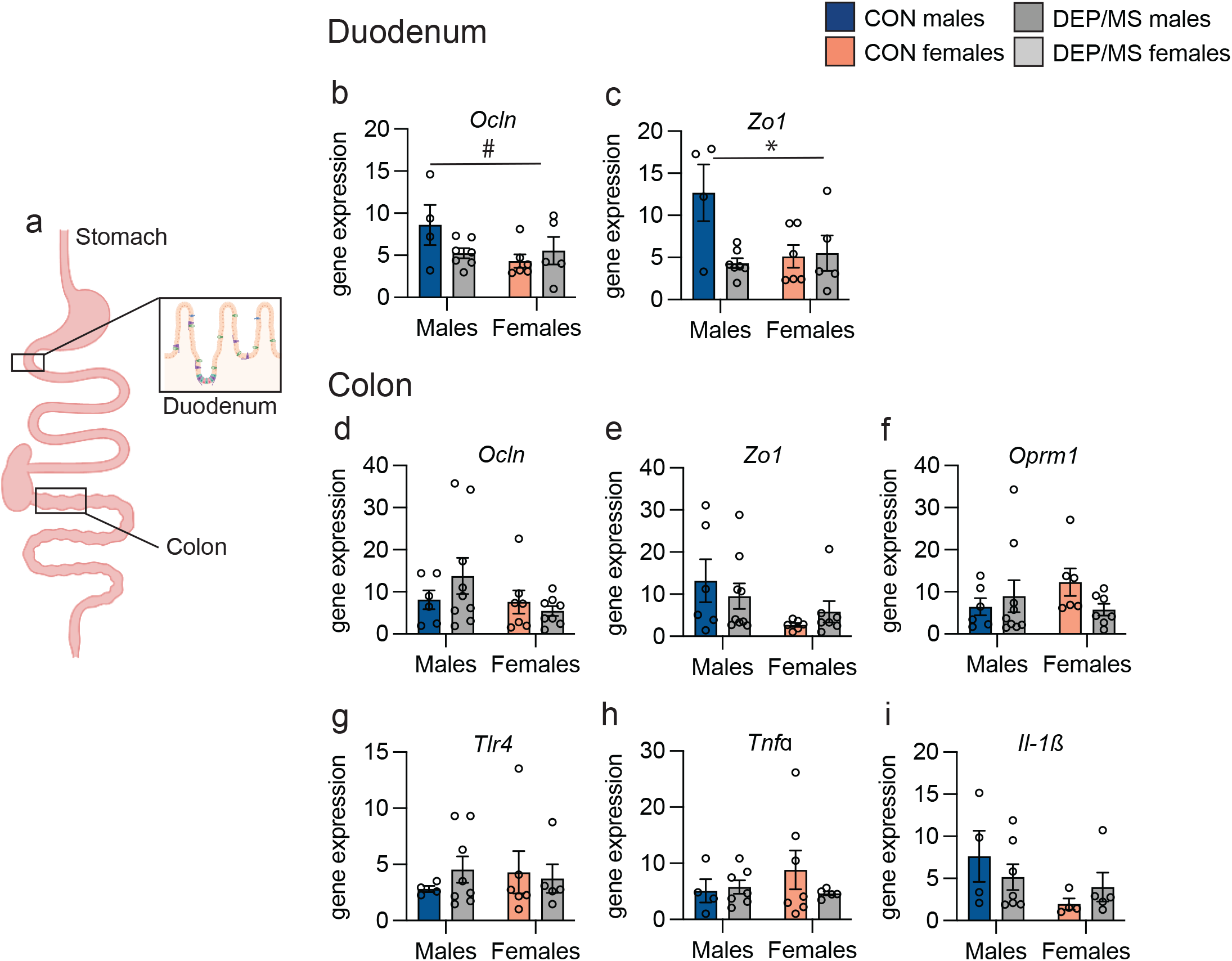
DEP/MS impacts gene expression in the duodenum but not the colon. **a**, Schematic depicting collection sites within the intestine for duodenum and colon samples. **b-c**, Treatment and sex tended to interact for *Ocln* (**b**) and significantly interacted for *Zo1* (**c**) mRNA in the duodenum - recapitulating findings in the ileum (N=4-9/group, 2-way ANOVA [treatment x sex], *Ocln*: treatment: p=0.09, *Zo1*: treatment: p=0.02). **d-i**, no significant differences in *Ocln* (**d**), *Zo1* (**e**), *Oprm1* (**f**), *Tlr4* (**g**), *TNFa* (**h**), or *ll-1ß*, (**i**) gene expression were observed in the colon following DEP/MS (N=4-9/group, 2-way ANOVA [treatment x sex]. Data represent mean +/- SEM *p<0.05, #p<0.1.

**Extended Figure 7.**
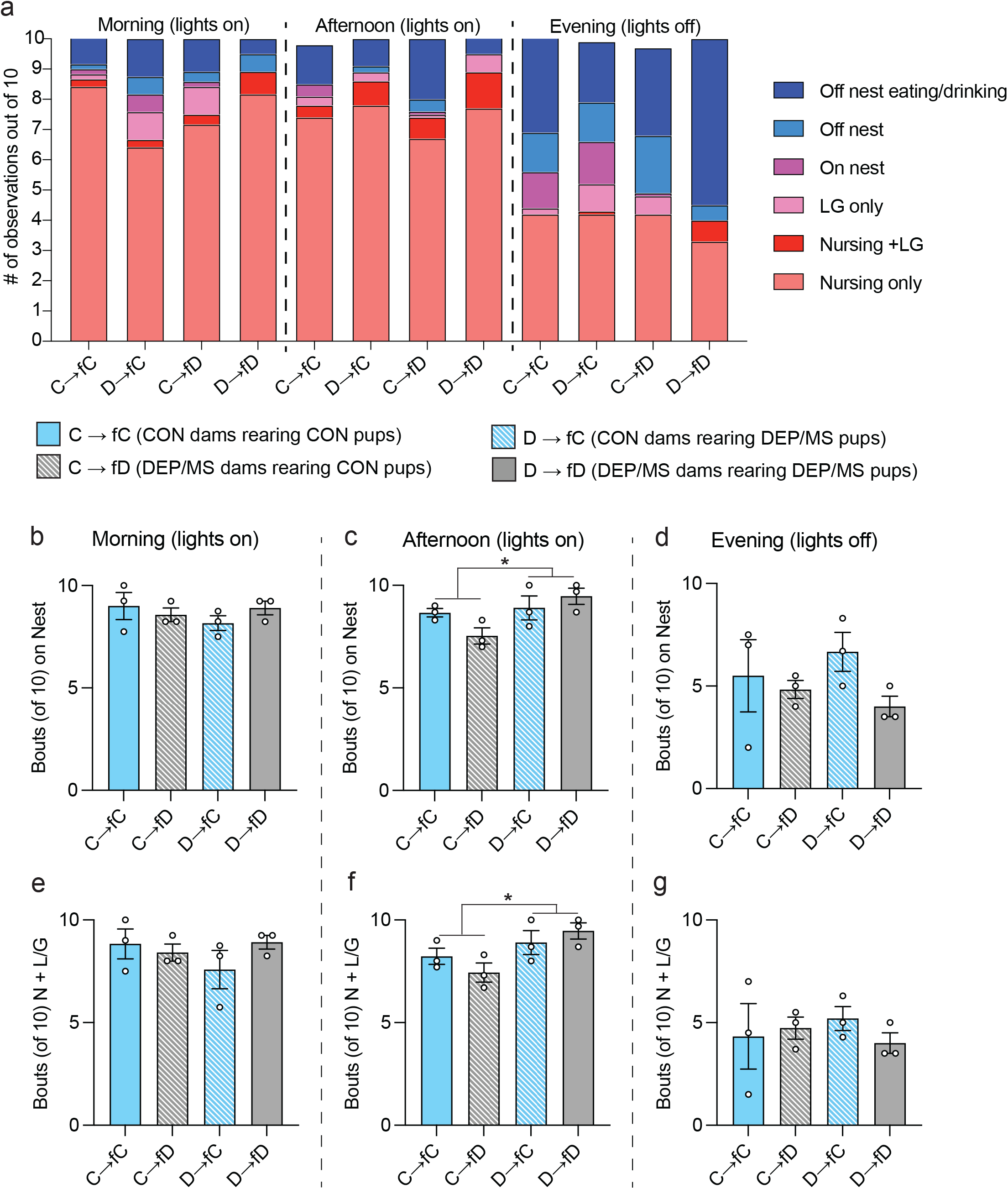
Expanded effects of DEP/MS on maternal behavior. **a**, Histogram of group means of maternal behaviors over the different times of day (averaged across the first 3 days of life). Maternal behaviors while on the nest (in warm colors) were categorized as ‘on nest’, ‘nursing (N)’, ‘licking/grooming (LG)’, nursing and licking/grooming (N + LG)’. Behaviors off the nest (cool colors) were categorized as ‘Off nest’ or ‘off nest eating/drinking’. No active maternal behaviors were observed off the nest. **b**, During the morning observation period, there was no significant effect of either dam or pup treatment condition on Bouts in Nest or Bouts of N+ LG (**e**, N=3/group, 2-way ANOVA (pup treatment x dam treatment), Bouts on Nest: pup treatment p=0.59, dam treatment p=0.72, interaction p=0.23, Bouts of N+LG: pup treatment p=0.58, dam treatment p=0.50, interaction p=0.21). **c**, In the afternoon observation period, there were significant effects of pup treatment condition on both Bouts in Nest and Bouts of N+ LG (**f**, N=3/group, 2-way ANOVA (pup treatment x dam treatment), Bouts on Nest: pup treatment p=0.03, dam treatment p=0.52, interaction p=0.08, Bouts of N+LG: pup treatment p=0.02, dam treatment p=0.81, interaction p=0.18). **d**, In the evening observation period, there were no significant effects of dam or pup treatment condition on Bouts in Nest or Bouts of N+ LG (**e**, N=3/group, 2-way ANOVA (pup treatment x dam treatment), Bouts on Nest: pup treatment p=0.88, dam treatment p=0.15, interaction p=0.37, Bouts of N+LG: pup treatment p=0.94, dam treatment p=0.68, interaction p=0.41).in **b-g**, dots represent animal averages across all 3 days analyzed. Data represent Mean +/- SEM, *p<0.05. CON: vehicle and control, DEP/MS: diesel exhaust particles/maternal stress.

**Extended Figure 8.**
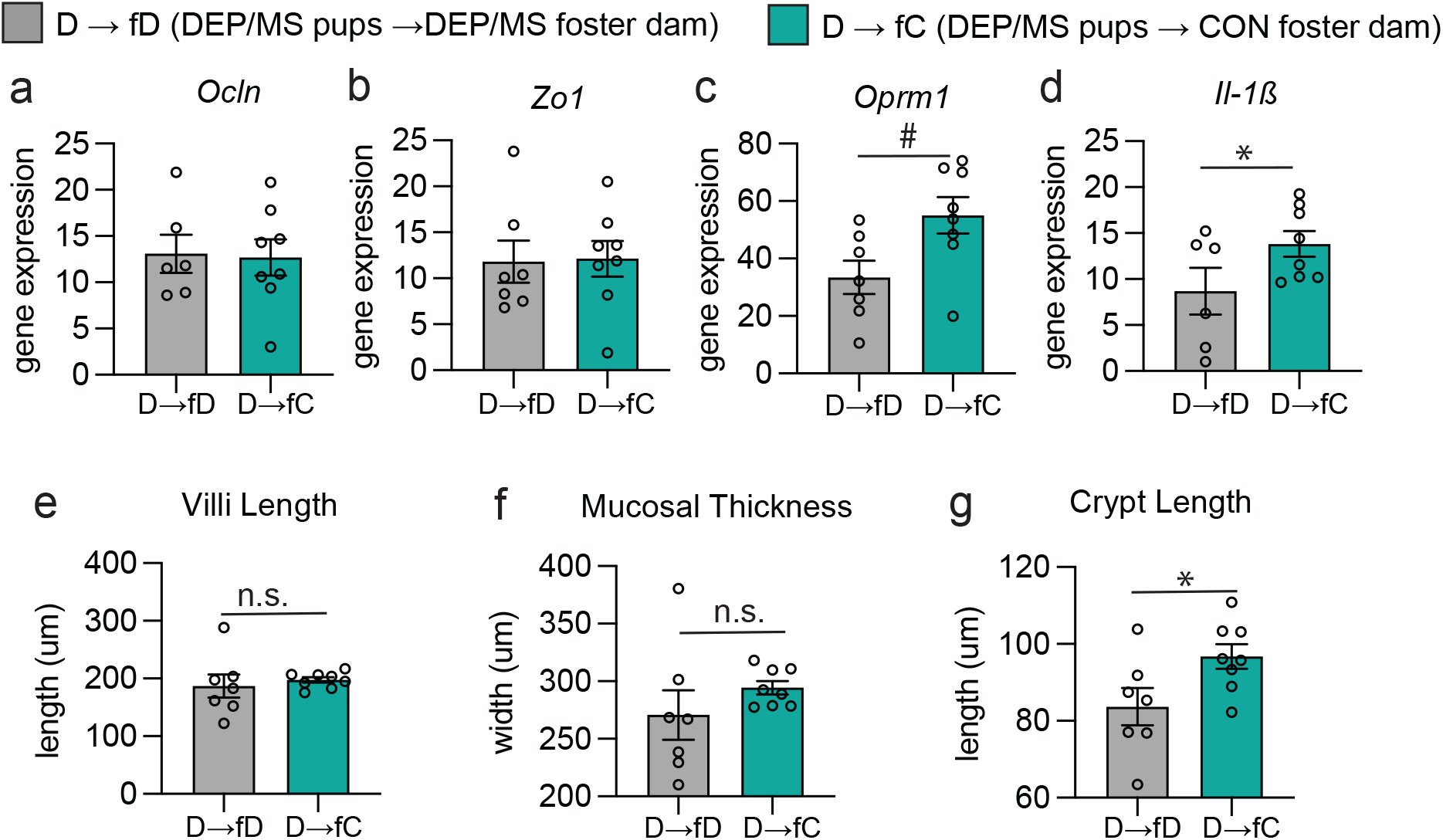
Effects of cross-fostering to a CON dam at birth on the intestinal epithelium in DEP/MS male offspring. **a-b**, Neither *Ocln* nor *Zo1* gene expression differed between DEP/MS pups reared by a DEP/MS or CON dam (N=6-8/group, un-paired t-test, *Ocln*: p=0.88, *Zo1*: p=0.92). **c-d**, Cross-fostering to a CON dam at birth tended to increase expression of *Oprm1* and significantly increased *Il-1ß* mRNAin DEP/MS male pups (N=6-8/group, un-paired t-test, *Oprm1*: p=0.08, *Il-1ß*: p=0.024). **e-f**, Neither villi length nor mucosal thickness differed between DEP/MS pups reared by a DEP/MS or CON dam. However, crypt length was increased in DEP/MS pups following fostering to a CON dam (N=7-8/group, un-paired t-test, villi length: p=0.60, mucosal thickness: p=0.28, crypt length: p=0.39, **g**). Data represent mean +/- SEM *p<0.05, # p<0.1, CON: Control, DEP/MS: diesel exhaust particle/maternal stress.

**Extended Figure 9.**
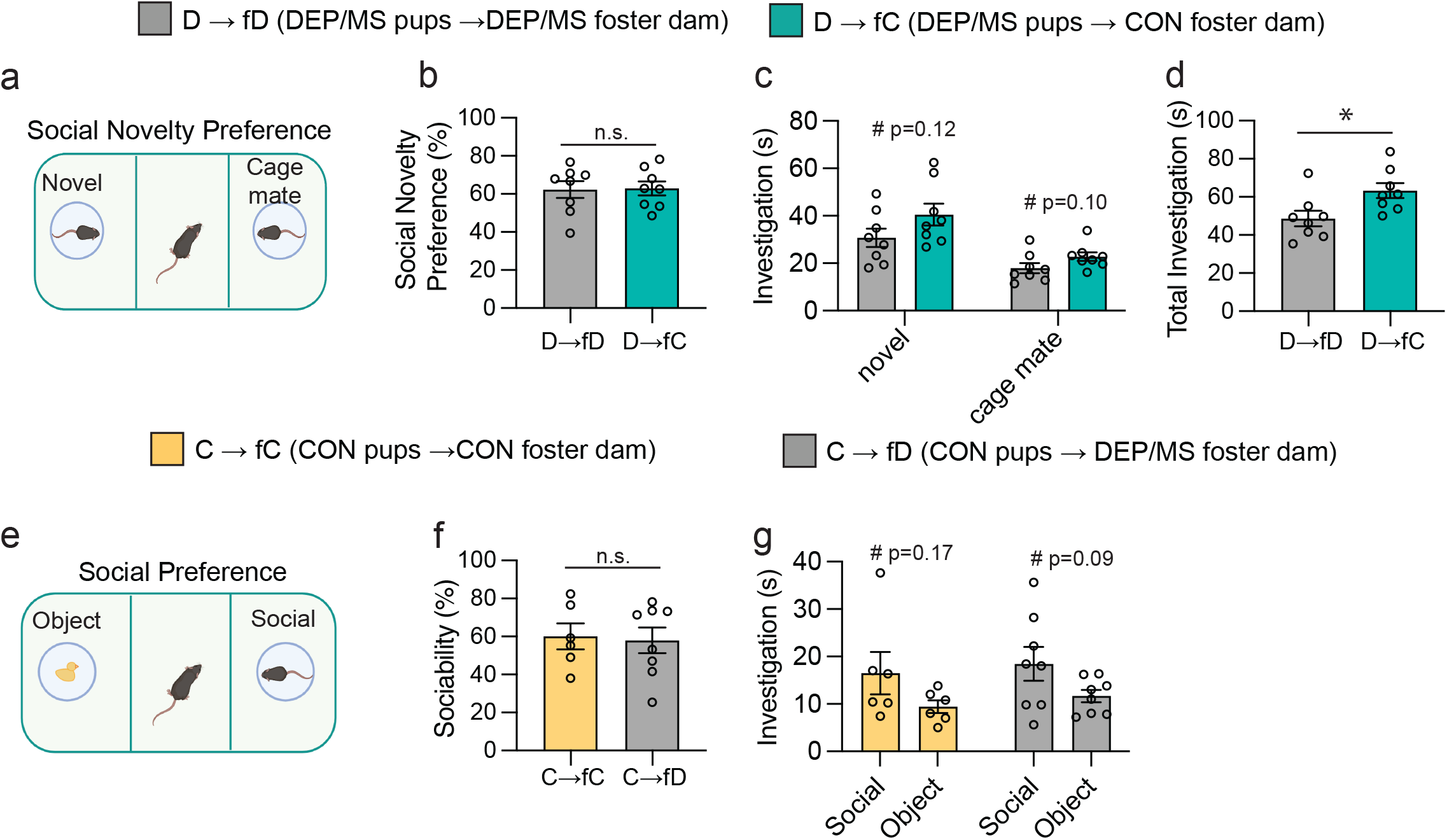
Extended effects of cross-fostering on social behavior. **a**, Representation of social novelty preference test. **b,** Cross-fostering to a CON dam at birth did not alter social novelty preference in DEP/MS pups (N=8/group, unpaired t-test [D → fD vs. D → fC], p=0.92). However, there was a trend towards higher novel and cage mate investigation in DEP/MS pups reared by CON dams (c, N=8/group, unpaired t-test [D → fD vs. D → fC], Novel Investigation: p=0.12, Cage mate Investigation: p=0.10) which drove an increase in total investigation time (**d**, N=8/group, unpaired t-test [D → fD vs. D → fC], p=0.02). **e**, Representation of social preference test. **f**, Cross-fostering of CON pups to a DEP/MS dam at birth did not induce social behavior deficits (N=6-8/group, unpaired t-test [C → fC vs. C → fD], p=0.83). **g**, Both CON and DEP/MS reared CON pups tended to spend more time interacting with a social stimulus as compared to an object (N=6-8/group, unpaired t-tests [Social vs. Object Investigation], C → fC: p=0.16, C → fD: p=0.097). Data represent mean +/- SEM *p<0.05, #p<0.15, CON: Control, DEP/MS: diesel exhaust particle/maternal stress.

